# The Histone Chaperone CAF-1 Sustains Myeloid Lineage Identity

**DOI:** 10.1101/2020.10.22.350447

**Authors:** Yiming Guo, Fei Ji, Jernej Murn, David Frankhouser, M Andres Blanco, Carmen Chiem, MiHyun Jang, Ruslan Sadreyev, Russel C. Rockne, David B. Sykes, Konrad Hochedlinger, Sihem Cheloufi

**Affiliations:** Department of Biochemistry, Stem Cell Center, University of California, Riverside, 900 University Ave. Boyce Hall 4411, Riverside, CA 92521-0129; Department of Molecular Biology, Massachusetts General Hospital, 185 Cambridge Street, Boston, MA 02114, USA; Department of Population Sciences and Department of Diabetes Complications & Metabolism, City of Hope, National Medical Center, Duarte, CA, United States; Department of Biomedical Sciences, School of Veterinary Medicine, University of Pennsylvania, Philadelphia, PA, United States; Department of Computational and Quantitative Medicine, Division of Mathematical Oncology, City of Hope National Medical Center, Duarte, CA, United States; Center for Regenerative Medicine, Massachusetts General Hospital, 185 Cambridge Street, Boston, MA 02114, USA; Department of Stem Cell and Regenerative Biology, Harvard University, Cambridge, MA 02138, USA; Harvard Stem Cell Institute, 1350 Massachusetts Avenue, Cambridge, MA 02138, USA; Cancer Center, Massachusetts General Hospital, 185 Cambridge Street, Boston, MA 02114, USA

## Abstract

During hematopoiesis, stem and progenitor cells become progressively restricted in their differentiation potential. This process is driven by lineage-specific transcription factors and is accompanied by dynamic changes in chromatin structure. The chromatin assembly factor complex CAF-1 is a key regulator of cellular plasticity in various cell lineages in different organisms. However, whether CAF-1 sustains lineage identity during normal homeostasis is unclear. To address this question, we investigated the role of CAF-1 in myeloid progenitor cells. CAF-1 suppression in myeloid progenitors triggered their rapid commitment but incomplete differentiation toward granulocyte, megakaryocyte, and erythrocyte lineages, resulting in a mixed cellular state. Through comparison with a canonical paradigm of directed terminal myeloid differentiation, we define changes in chromatin accessibility that underlie a unique transcriptome of the aberrantly matured CAF-1 deficient cells. We further identify C/EBPα and ELF1 as key transcription factors whose control of myeloid lineage commitment is kept in check by CAF-1. These findings shed new light on molecular underpinnings of hematopoiesis and suggest that manipulation of chromatin accessibility through modulating CAF-1 levels may provide a powerful strategy for controlled differentiation of blood cells.

## Introduction

Hematopoiesis involves sequential commitment of self-renewing hematopoietic stem cells (HSCs) to fully mature specialized blood cell types(Laurenti and Gottgens, 2018). During this process, HSCs become progressively more restricted towards megakaryocytes/platelets, erythroid, myeloid or lymphoid lineages by a stepwise transition through progenitor cell states. Recent single-cell transcriptome analyses of bone marrow suggest that lineage commitment is heterogeneous and deviates from the largely marker-based hierarchical differentiation model of hematopoiesis(Carrelha et al., 2018, Jacobsen and Nerlov, 2019, Laurenti and Gottgens, 2018, Liggett and Sankaran, 2020, Paul et al., 2016, Zhang et al., 2018). This signifies the need to understand the molecular mechanisms that sustain the identity of stem and progenitor cells and restrict their commitment to a specific lineage during differentiation.

Lineage specification during hematopoiesis is tightly controlled by transcription factors. In the myeloid lineage, the CCAAT/enhancer-binding protein (C/EBP) family members play major roles in commitment toward granulocytes and macrophages, while GATA1, KLF1, and GFI1B have been described to govern erythrocyte and megakaryocyte lineage commitment(Rosenbauer and Tenen, 2007). Notably, some of these transcription factors, such as C/EBPα and GATA1, are sufficient to drive transdifferentiation when ectopically expressed in a different blood cell lineage(Xie et al., 2004, Heyworth et al., 2002). However, the mechanisms, including chromatin-regulatory processes, that initiate early myeloid lineage commitment remain poorly defined.

During the dynamic process of cellular differentiation, chromatin remodeling typically precedes transcriptional regulation(Atlasi and Stunnenberg, 2017). Therefore, it is important to dissect how altered chromatin accessibility sets the stage for the activity key transcription factors as drivers of cell lineage specification. Of the many types of molecules implicated in the control of chromatin accessibility, histone chaperones act broadly by catalyzing nucleosome assembly during DNA replication, gene transcription, and DNA repair(Hammond et al., 2017). The chromatin assembly factor complex CAF-1 assembles nucleosomes in a DNA replication-dependent manner. We have previously identified two subunits of the CAF-1 complex, Chaf1a and Chaf1b, as key regulators of cell identity maintenance in different paradigms of transcription factor-driven cellular reprogramming and direct lineage conversions(Cheloufi et al., 2015, Cheloufi and Hochedlinger, 2017). Mechanistically, CAF-1 blocks the binding of ectopically expressed transcription factors by maintaining a closed chromatin state at target loci. Consistent with this observation, loss of CAF-1 enhances transcription factor-driven reprogramming of somatic to pluripotent stem cells and direct lineage conversion of pre-B cells into macrophages and that of fibroblasts into neurons. Given the effect of the CAF-1 complex on chromatin accessibility in these and other cellular paradigms(Cheloufi et al., 2015, Cheloufi and Hochedlinger, 2017, Ishiuchi et al., 2015), CAF-1 has been viewed as a general stabilizer of cell identity that prevents cells from adopting open chromatin state characteristic of immature cells. However, studies of CAF-1 function in mammalian tissue homeostasis have been aggravated due to its requirement during DNA replication and hence its essential role in cell proliferation and organismal development(Houlard et al., 2006, Volk et al., 2018, Cheloufi and Hochedlinger, 2017). Recent studies of embryonic stem cell (ESC) differentiation and T cell development uncovered additional roles of CAF-1 in cooperating with chromatin-modifying enzymes, such as the polycomb repressive complex 2 (PRC2), histone deacetylases (HDAC1/2), the histone demethylase LSD1 and DNA methyltransferase(Cheng et al., 2019, Ng et al., 2019). However, how these activities of CAF-1 might be coupled to the action of transcription factors and regulation of cellular identity in normal homeostasis remains to be determined. Here we combine a controllable myeloid differentiation system with inducible perturbation of CAF-1 to investigate its role in sustaining myeloid lineage integrity and identify the early transcriptional events that control blood lineage fidelity.

## Results

### Loss of CAF-1 relieves the differentiation block in myeloid progenitors

During normal homeostasis, granulocyte-macrophage progenitors (GMPs) are a transient and highly proliferative blood cell population committed to differentiate into neutrophils and macrophages(Laurenti and Gottgens, 2018). GMPs can be propagated in culture as a stable self-renewing population that retains a normal karyotype and normal differentiation potential through overexpression of the Homeobox A9 transcription factor HOXA9 (hereafter Hoxa9-GMPs)(Wang et al., 2006, Sykes et al., 2016). In this system, the hormone-binding domain of the estrogen receptor (ER) is fused to the N-terminus of Hoxa9 to allow for estradiol-regulated nuclear localization and thus transcriptional activity of the ER-Hoxa9 fusion protein. Withdrawal of estradiol from culture triggers homogenous differentiation of Hoxa9-GMPs into neutrophils within 4 days (Figure. 1a). The Hoxa9-GMPs harbor a lysozyme-GFP reporter transgene (Lys-GFP) to allow for monitoring of cell differentiation towards granulocytes(Faust et al., 2000). Importantly, Hoxa9 inactivation through estrogen withdrawal mirrors the natural differentiation of primary mouse and human GMPs(Sykes et al., 2016).

**Figure 1:**
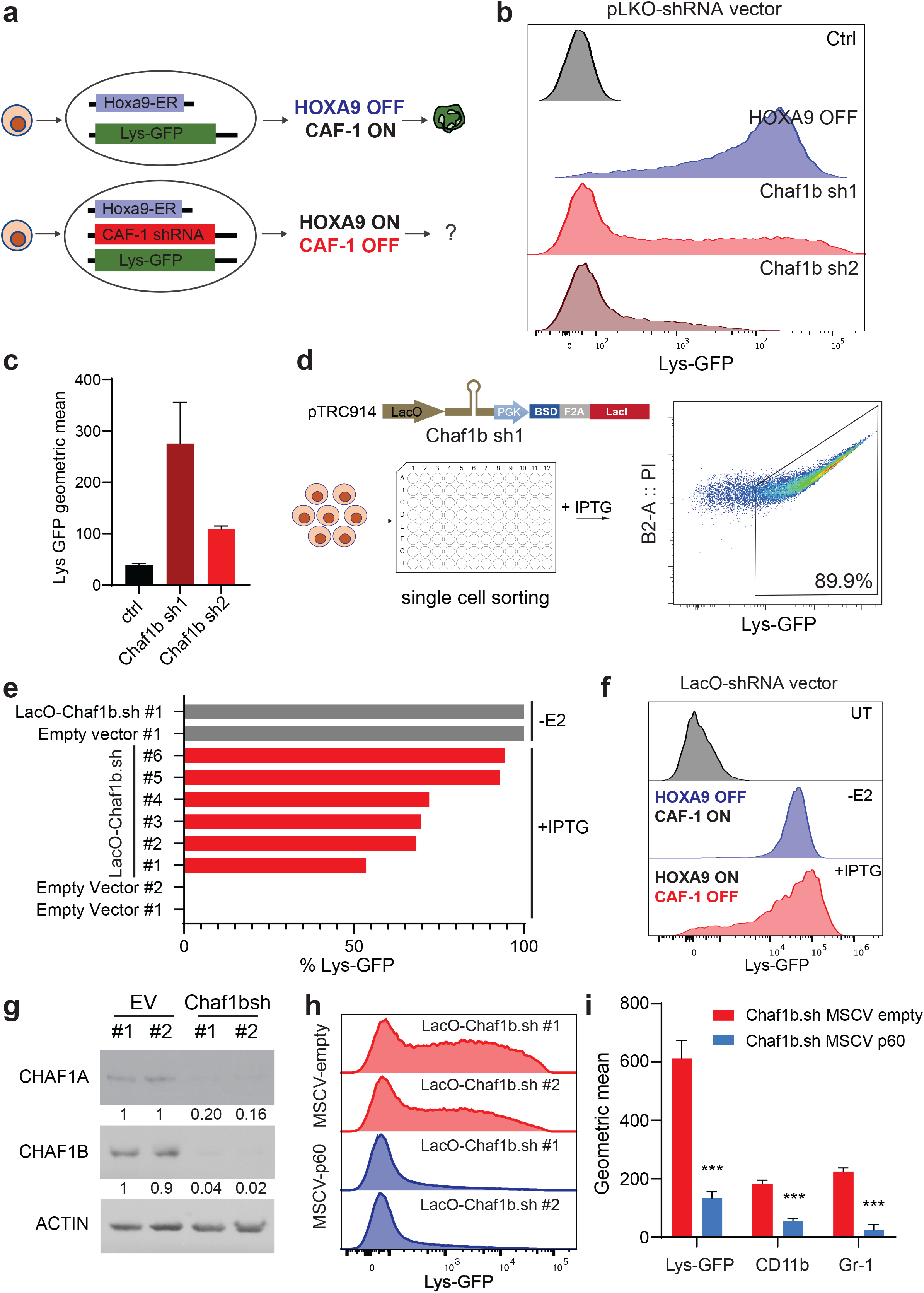
Loss of CAF-1 relieves the differentiation block in myeloid progenitors. **a,** Schematic of GMP differentiation system and CAF-1 perturbation strategy. GMPs are lock in undifferentiated state with Hoxa9-ER transgene (blue). Lys-GFP (green) is a reporter of granulocytes differentiations. Chaf1b shRNAs lentiviral vectors (red) are introduced in the presence of active HOXA9. **b,** Flow cytometric analysis of Lys-GFP expression upon constitutive loss of Chaf1b in Hoxa9-GMPs using two independent shRNAs. **c,** Quantification of data shown in **b**. Values are the mean from two independent experiments. **d,** IPTG RNAi inducible system for Chaf1b suppression and strategy for establishment of monoclonal inducible cell lines. **e,** Flow cytometric analysis of Lys-GFP expression in monoclonal cell line selection representing variegated strength of differentiation phenotypes in 6 independent subclones upon IPTG administration. Hoxa9 inactivation upon β-Estradiol withdrawal (-E2) is used as a control. **f,** Representative flow cytometric analysis of Lys-GFP expression in the best subclone #1. (UT, untreated). **g,** Western blot analysis confirming shRNA suppression of CHAF1B protein and subsequent degradation of the CHAF1A subunit in the two best Chaf1b shRNA inducible sub-clones (#1&#2) compared to control clones treated with IPTG. **h,** Differentiation rescue using RNAi resistant Chaf1b cDNA represented by flow cytometric analysis of Lys-GFP expression in IPTG inducible Chaf1b shRNA monoclonal cell lines transduced with either Chaf1b cDNA or empty vector control. **i,** Quantification of flow cytometry data shown in **h** and additional granulocytes differentiation markers (CD11b and Gr-1). Values are the mean from biological triplicates; error bars indicate standard deviation (***P < 0.001)

To test whether the CAF-1 complex maintains GMP identity, we used shRNAs targeting its Chaf1b subunit. Two independent Chaf1b shRNAs resulted in activation of the Lys-GFP reporter bypassing the differentiation blockade by Hoxa9 (Figure 1b-c). To allow for tunable repression of Chaf1b we cloned the Chaf1b shRNAs into a lactose operator (Lac-O) inducible system that allows for IPTG regulated expression of the shRNA transgene. Although the lac-O-driven Chaf1b shRNA-transduced polyclonal population showed moderate level of differentiation compared to the constitutively driven shRNA, sub-clones of cells showed up to 90% activation of the Lys-GFP reporter, mirroring the effect of Hoxa9 inactivation (Figure 1d-f). The inducible Chaf1b shRNA reduced protein levels of CHAF1A and CHAF1B proteins (Figure 1g), in line with the previously observed co-dependent stability of the two CAF-1 subunits. We therefore refer to Chaf1b knockdown as CAF-1 OFF and Hoxa9 inactivation as HOXA9 OFF. Importantly, overexpression of an RNAi-resistant Chaf1b cDNA in the Lac-O-Chaf1b shRNA-expressing clones rescued the differentiation phenotype, as judged by low expression of Lys-GFP and two additional granulocyte differentiation markers, CD11b and Gr-1 (Figure 1h, i).

### CAF-1 inhibition induces rapid but partial differentiation of Hoxa9-GMPs

To gain insight into the dynamics of differentiation mediated by CAF-1 inhibition or Hoxa9 inactivation, we performed a series of experiments to survey the early differentiation events and the point of commitment. We detected activation of neutrophil differentiation markers (Lys-GFP, CD11b and Gr-1) as early as 48hrs in both CAF-1 OFF and Hoxa9 OFF conditions (Figure. 2a-b & Figure2-figure supplement 1a-d). Consistently, CAF-1 depletion resulted in similar growth arrest compared to Hoxa9 inactivation (Figure2-figure supplement 1e). We note that the dynamics and strengths of the observed phenotypes corresponded with reduced protein levels of the CHAF1B and CHAF1A subunits, which were apparent in both CAF-1 OFF and Hoxa9 OFF conditions (Figure 2c). To investigate the speed with which cells commit to a stable differentiated state, we pulsed cells in a time-course experiment with either IPTG (CAF-1 OFF) or estradiol withdrawal (Hoxa9 OFF), followed by a chase period before assessment of cell differentiation based on the Lys-GFP reporter (Figure. 2d). Markedly, within 48hrs of either pulse, the differentiated Hoxa9-GMPs did not revert during the ensuring 96hrs chase period (Figure. 2d). Moreover, the proportion of differentiated cells increased with longer pulse periods, reaching a large majority at 96hrs in both treatments (> 90% of differentiated Hoxa9-GMPs). Together, these results indicate that loss of CAF-1 releases the Hoxa9-mediated differentiation block as early as 48hrs reaching a maximum effect within 96hrs mirroring the effect of normal GMP differentiation.

**Figure 2:**
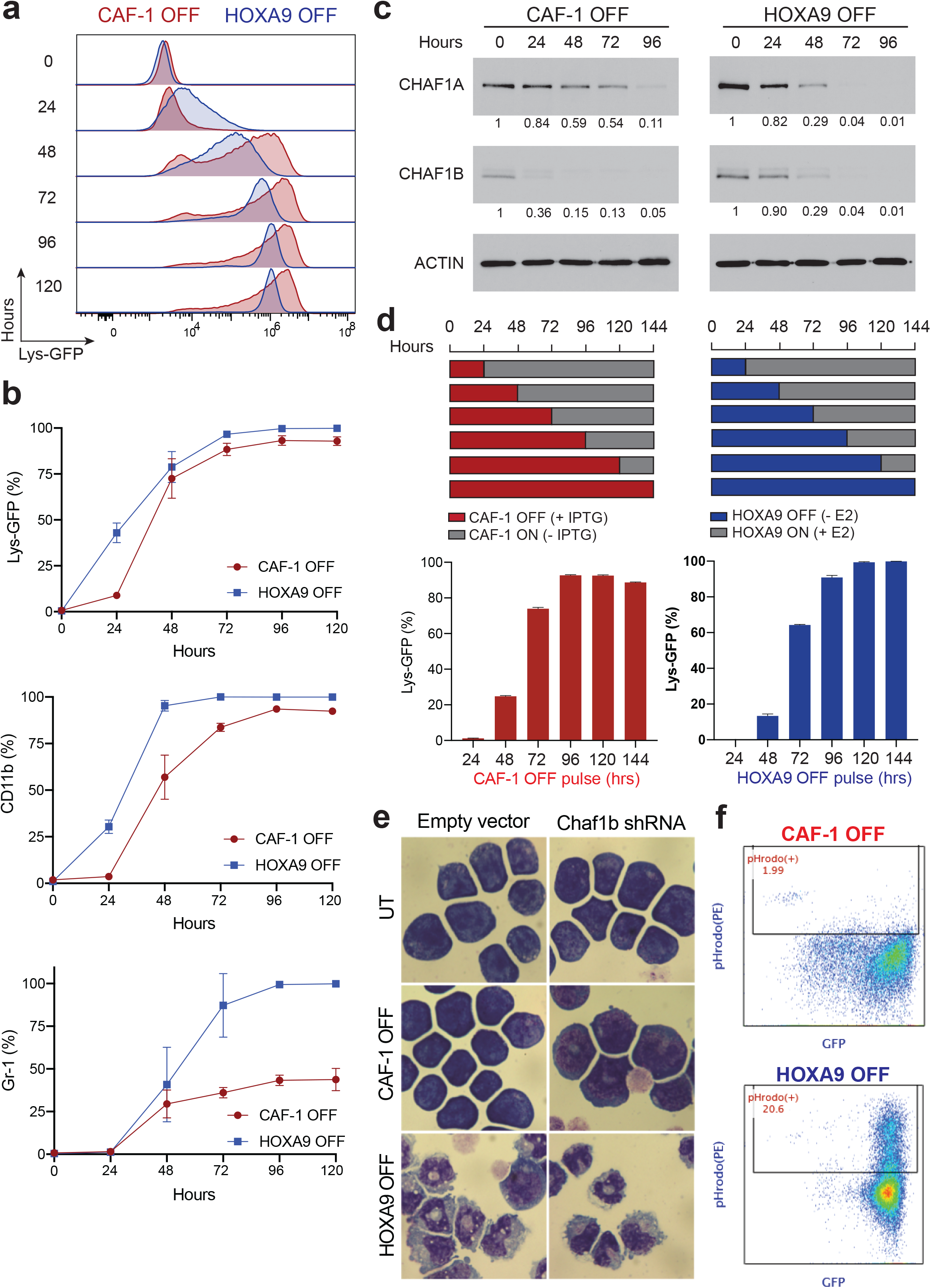
CAF-1 inhibition induces rapid but partial differentiation of Hoxa9-GMPs. **a,** Time course analysis of GMP differentiation during 5 days pulse period of IPTG (CAF-1 OFF) or -E2 (HOXA9 OFF) using flow cytometric analysis of Lys-GFP expression. **b,** Time course analysis of a set of granulocyte marker activation using flow cytometric analysis of Lys-GFP, CD11b and Gr-1 expression in CAF-1 OFF versus HOXA9 OFF GMPs. Representative example of Lys-GFP positive cells is shown in the histogram in **a**. **c,** Western blot time course analysis during 4 days pulse period depicting the shRNA suppression of CHAF1B protein and subsequent degradation of the CHAF1A subunit in CAF-1 OFF cells (left panel). CHAF1B and CHAF1A subunits are also naturally reduced in HOXA9 OFF cells (right panel). **d**, Pulse-Chase of either CAF-1 OFF (left plot) or HOXA9 OFF (right plot) to determine the point of commitment to granulocyte fate using flow cytometric analysis of Lys-GFP expression. X axes represents the pulse period followed by the corresponding chase period represented in the top schematic panel. CAF-1 OFF and HOXA9 OFF cells retain Lys-GFP activation after only 48hrs pulse and 5days chase. **e,** Morphologic analysis of CAF-1 OFF and HOXA9 OFF cells using Wright-Giemsa staining at 96hrs in a control clone and Chaf1b shRNA IPTG inducible clones. Shown are representative images from the analysis of two independent controls (left) and Chaf1b shRNA expressing subclones (right). UT (untreated cells with GMP morphology). **f,** Phagocytosis flow cytometric assay of CAF-1 OFF and HOXA9 OFF at 96hrs measuring the engulfment of fluorescently labeled bacterial particles.

To further investigate the fate of CAF-1 depleted Hoxa9-GMPs, we performed morphologic cyto-spin analysis and phagocytosis assays. Despite the growth arrest and marked induction of neutrophil cell surface markers, CAF-1 suppression did not induce the fully-mature neutrophil morphology or the ability of cells to engulf fluorescently labeled bacterial particles, traits seen in the Hoxa9 OFF condition (Figure. 2e-f and Figure2-figure supplement 1f). Taken together, these observations suggest that acute CAF-1 loss in Hoxa9-GMPs triggers rapid but partial granulocyte differentiation.

### Co-activation of multilineage genes upon CAF-1 suppression in Hoxa9-GMPs

To better characterize the identity of CAF-1 depleted Hoxa9-GMPs and considering a possibility that these cells might present a heterogeneous population, we conducted single cell transcriptome analysis at 48h and 96h inductions of CAF-1 OFF and Hoxa9 OFF states (Figure3-figure supplement 1a-c). Using Uniform Manifold Approximation and Projection (UMAP) analysis(Becht et al., 2018), we found diverging cell clusters clearly distinguishing both cellular states, a trend that became progressively more apparent with increasing time of differentiation (Figure. 3a). Fine-resolution unsupervised clustering of differentially expressed genes between single cells confirmed this finding and identified several gene sets distinguishing the two differentiation trajectories (Figure. 3a, b). For instance, the top differentially regulated genes with relative enrichment in the Hoxa9 OFF state included granulocyte maturation markers s100a8, s100a9 and Ltf. In contrast, cells in the CAF-1 OFF state exhibited markedly higher expression of (1) genes broadly associated with hematopoietic progenitors (e.g, Cd34, Prtn3), (2) histone variant genes (H3f3b, Cenpa, Hist1h2ap and H1fx), and (3) the platelet factor 4 (Pf4) transcription factor, a specific marker of megakaryocyte progenitor (MegP) cells. Moreover, several early granulocyte-associated genes, including Fcnb, Elane and Ms4a3, showed, as expected, only transient expression upon Hoxa9 inactivation, but a sustained upregulation in the CAF-1 OFF condition (Figure. 3b). These clustering patterns are consistent with the above observation that CAF-1 depletion leads to incomplete differentiation of myeloid progenitors.

**Figure 3:**
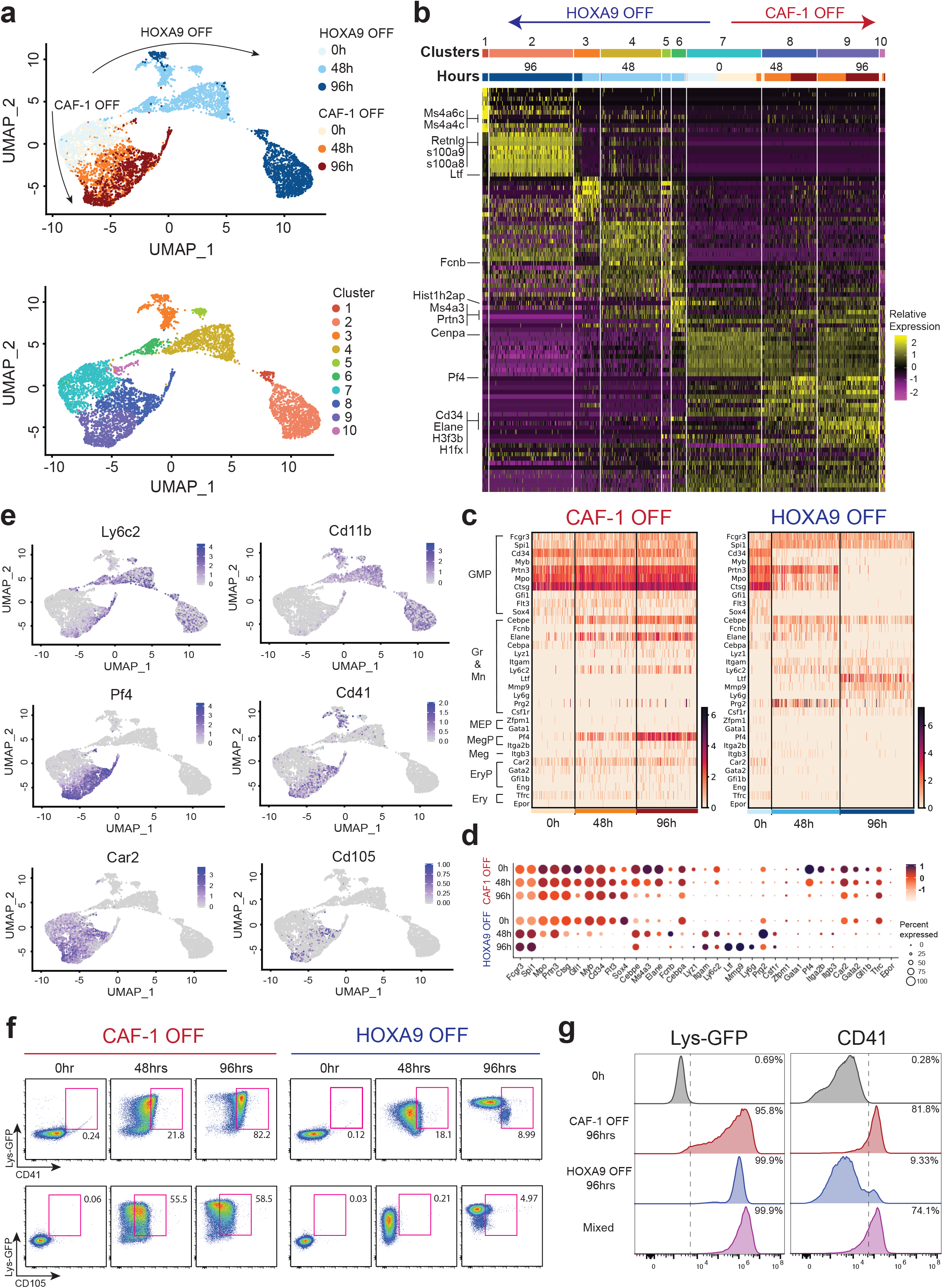
Co-activation of multilineage genes upon CAF-1 suppression in Hoxa9-GMPs. **a,** UMAP displaying 10X Chromium scRNA-Seq data, colored by time points (upper panel) and sub-clusters (lower panel). Clustering performed with SLM algorithm using PC1-10. **b,** Heatmap displaying the top 10 enriched genes in each sub-cluster (shown in **a**, lower panel) ordered by time points and conditions. Select DEGs from different categories discussed in the text are highlighted by insets to the left of the heatmap. **c,** Heatmap displaying gene expression pattern of a curated set of lineage specific genes (surface markers and transcription factors) at different time points in CAF-1 OFF versus HOXA9 OFF cells. **d**, Bubble plots reflecting the lineage markers shown in **c** in CAF-1 OFF versus HOXA9 OFF cells. The color of the dots shows the relative expression level and the size of the dots represents the percentage of cells that express the corresponding gene. **e,** UMAP displaying the expression of select markers that characterize different lineages: Ly6c2 and CD11b for neutrophils, Pf4 and Cd41 for megakaryocyte progenitors and Car2 and Cd105 for erythrocyte progenitors. **f**, Flow cytometric analysis of Lys-GFP, CD41 and CD105 expression at 0, 48, and 96 hrs in CAF-1 OFF versus HOXA9 OFF cells reflecting the mixed cellular state of CAF-1 OFF cells only. Lys-GFP and CD41 or CD105 double positive cells are gated. **g**, Representative flow cytometric analysis of Lys-GFP and CD41 activation upon subsequent (IPTG and -E2) pulse treatments. A representative longest pulse is shown here with initial CAF-1 OFF for 96hrs followed by HOXA9 OFF for 96hrs showing a modest rescue of the mixed cellular state when HOXA9 is subsequently inactivated in CAF-1 OFF GMPs. Individual CAF-1 OFF and HOXA9 OFF are shown as controls. Fine-tuned incremental pulse analysis is shown in Figure 3-figure supplement 2.

To resolve whether CAF-1 OFF cells are individually locked into a mixed cellular state or exhibit heterogeneity only as a population, we scrutinized a curated set of hematopoietic lineage genes, including cell surface markers and transcription factors. CAF-1 OFF cells retained progenitor-associated genes Cd34, Prtn3, and Flt3 indicating an incomplete exit of these cells from the progenitor stage (Figure. 3c-d). Interestingly, we also detected a marked upregulation of erythrocyte progenitor (EryP) markers (Car2, Gata2, Cd105, Gfi1b) and MegP markers (Pf4, Itga2b) but no noticeable induction of lymphoid lineage-specific genes (Figure. 3c-e and data not shown). We found that expression of the distinct lineage-specific markers was not limited to different subsets of cells but that the majority of the cells co-expressed several of these genes, indicating a mixed cellular state (Figure. 3c-e and Figure3-figure supplement 1d). To check whether these transcript-level observations might be manifested on the protein level, we assayed cell surface expression dynamics of the MegP marker Itga2b (CD41) and EryP marker CD105 in each differentiation condition. Consistent with the RNA-seq analysis, we found co-expression of both markers along with Lys-GFP uniquely upon CAF-1 suppression (Figure. 3f & Figure3-figure supplement 1 e-f). The mixed cellular state of CAF-1 OFF cells was recapitulated using an independent inducible sub-clone, which further pointed to a correlation between CAF-1 dosage and the strength of co-induction of the multilineage markers (Figure3-figure supplement 1g-h). The pervasive co-expression of erythrocytes and megakaryocytes genes in conjunction with the activation of myeloid differentiation genes and the retention of myeloid progenitor markers upon CAF-1 suppression in GMPs suggests their incomplete differentiation and a mixed cellular state.

To test whether the persistent activation of Hoxa9 in CAF-1 OFF cells could account for the cells expressing the diverse lineage markers, we inactivated Hoxa9 following incremental pulse depletions of CAF-1 up to 96hrs (Figure3-figure supplement 2a). This resulted in augmented upregulation of neutrophil marker activation (~100% Lys-GFP; Figure. 3g & Figure3-figure supplement 2b-c) and a modest downregulation of CD41 regardless of the length of the CAF-1 OFF pulse (Figure. 3g and Figure3-figure supplement 2d-e). These results suggest that loss of CAF-1 is the predominant cause of the Hoxa9-GMPs mixed cellular state.

### CAF-1 inhibits multiple transcriptional programs to block differentiation of Hoxa9-GMPs

To identify transcription regulators sustaining the mixed cellular state upon CAF-1 suppression in GMPs, we argued that putative changes in chromatin accessibility resulting from depletion of CAF-1 could provide a critical clue, in line with histone deposition as the primary function of the CAF-1 complex(Smith and Stillman, 1989). To capture potentially causative changes, we performed RNA-seq and ATAC-seq analyses at 48 hours, which we considered an early time point when cells readily initiate differentiation (Figure. 2a & d). As could be expected, an overwhelming majority (87.3%) of all differentially expressed genes (DEGs) in CAF-1 OFF cells were upregulated, in agreement with a gross increase in chromatin accessibility (66.1% of all changes showed increased accessibility; Figure. 4a-d). This contrasted with Hoxa9 OFF cells, where a substantially smaller majority of DEGs were upregulated (59.5%) and where chromatin accessibility showed little net change (50.7% gained versus 49.3% lost peaks; Figure. 4a-d). In addition, significantly larger totals numbers of DEGs (6.8-fold more) and changes in chromatin accessibility (2.2-fold more) were detected in Hoxa9 OFF compared to CAF-1 OFF conditions (Figure. 4a-b & Figure4-figure supplement 1a-b). These data support the disparate differentiation trajectories and the resulting cellular phenotypes induced by loss of CAF-1 or Hoxa9 (Figures. 2 and 3). The chromatin rearrangement and transcriptional program controlled by CAF-1 may, based on these data and the common suppression of the CAF-1 complex under normal GMP differentiation (Figure. 2c), present only a fraction of the much larger Hoxa9-regulated transcriptional network that supports terminal differentiation to neutrophils. Nevertheless, the contribution via loss of CAF-1 alone may be critical, as indicated by gene set enrichment analysis (GSEA)(Subramanian et al., 2005) and EnrichR(Kuleshov et al., 2016) analyses, which point to globally significant induction of genes linked to myeloid differentiation in both conditions (Figure4-figure supplement 1c, d).

**Figure 4:**
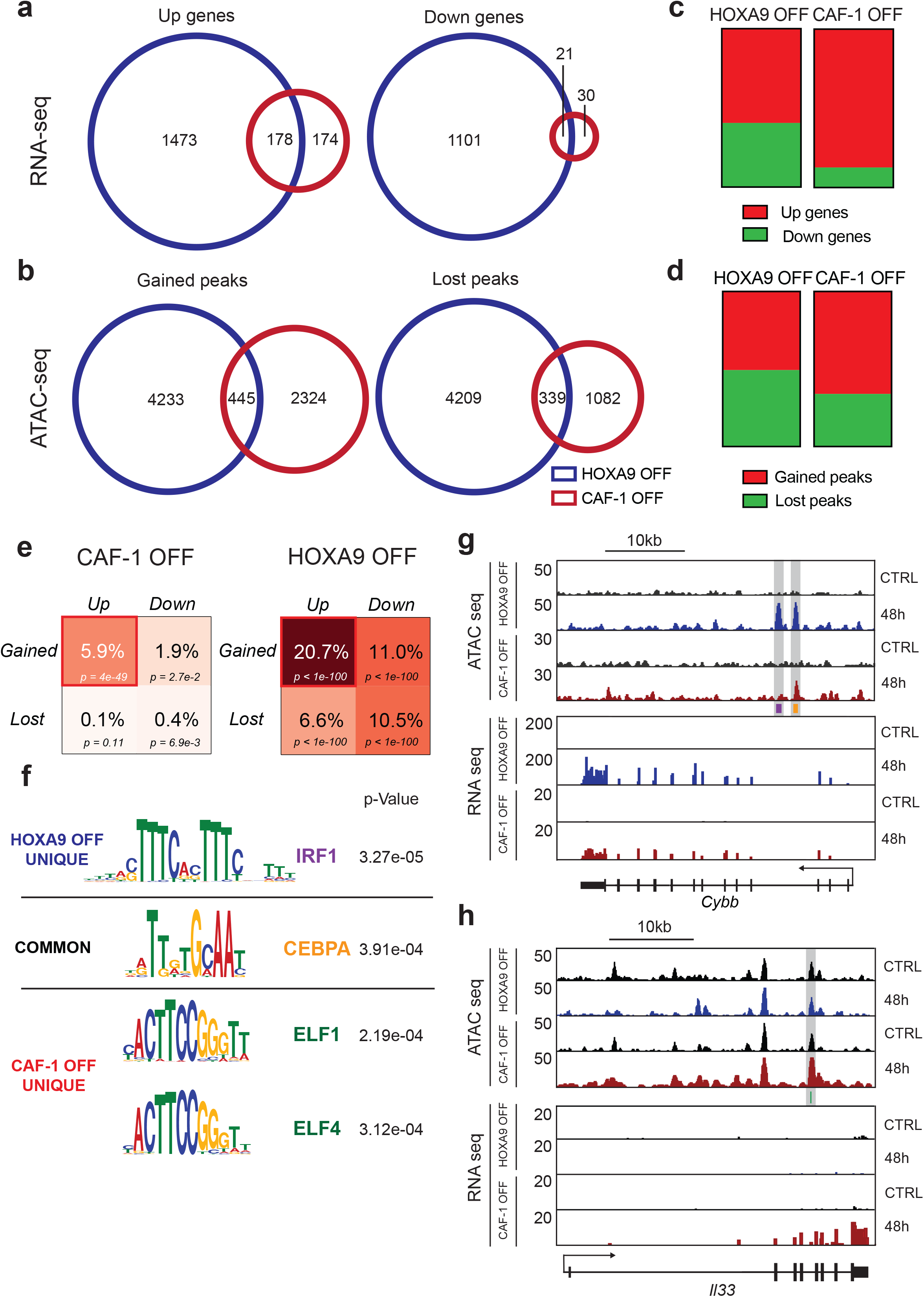
CAF-1 inhibits multiple transcriptional programs to block differentiation of GMPs. **a,b** Venn diagrams depicting common and unique DEGs (top panel) and differentially accessible ATAC-seq peaks (bottom panel) in CAF-1 OFF versus HOXA9 OFF cells at 48hrs. Upregulated genes and gained ATAC-seq peaks (left panels). Downregulated genes and lost ATAC-seq peaks (right panel). **c, d** Bar graph showing the fractions of differentially expressed genes and differentially accessible ATAC-seq peaks in CAF-1 OFF versus HOXA9 OFF GMPs at 48hrs. **e**, Fractions of differentially accessible ATAC-seq peaks in CAF-1 OFF versus HOXA9 OFF GMPs that are within 100KB distance from differentially expressed genes represented as a matrix comparing gain versus loss of chromatin accessibility and how it correlates with upregulation versus downregulation of neighboring genes. **f,** Transcription factor motifs enriched from gained ATAC-seq peaks that are common or unique between CAF-1 OFF and HOXA9 OFF conditions. **g, h** Representative examples of correlation shown in **f.** represented by genome tracks of ATAC-seq and RNA-seq peaks. **g,** example of a common target gene Cybb neighboring CEBPα (orange) and IRF1 (purple) predicted binding sites. **h**, example of a CAF-1 OFF unique target gene Il33 neighboring ELF1 predicted binding site (green).

Increased chromatin accessibility resulting from depletion of CAF-1 may promote chromatin associations of distinct transcription factors to drive the establishment of the observed mixed cellular state (Figure. 3). To identify such factors, we looked for enriched DNA motifs as potential transcription factor binding sites in the gained ATAC-seq peaks in CAF-1 OFF cells and compared them with the Hoxa9 OFF condition. Remarkably, a large percentage of gained sites in both conditions are within 100Kb of upregulated genes in both conditions (Figure. 4e). Multiple expression motifs for motif elicitation (MEME) (Machanick and Bailey, 2011) predictions identified C/EBPα binding motif as the most significantly enriched motif in the commonly gained accessible regions in both conditions (Figure. 4f). This is consistent with the known role of C/EBPα as a master regulator of GMP differentiation(Pundhir et al., 2018, Rosenbauer and Tenen, 2007). Given its expression profile and the requirement of C/EBPα during the early stages of myeloid differentiation (Figure4-figure supplement 2a-c), these results suggest that depletion of CAF-1 or Hoxa9 opens up chromatin regions that become accessible to C/EBPα, which in turn induces transcription of myeloid-specific genes. Indeed, we found several granulocytes specific genes in the vicinity of the gained C/EBPα target sites to be upregulated in both conditions (Figure. 4g and Figure4-figure supplement 2d-f). A similar search for DNA motifs enriched in chromatin regions that become accessible specifically upon CAF-1 depletion identified a sequence recognized by the E74 Like ETS transcription factors ELF1 and ELF4 (Figure. 4f), which are known primarily for their involvement in regulating immune responses(Gallant and Gilkeson, 2006). Elf1 in particular is more broadly expressed during hematopoiesis compared to C/EBPα (Figure4-figure supplement 2c) and has been proposed to regulate erythroid differentiation as well as development of natural killer cells and T cells(Suico et al., 2017). Markedly, we find potential Elf1 binding sites near genes upregulated only in the CAF-1 OFF condition (Figure. 4h and Figure4-figure supplement 2g). These observations suggest a contribution from the Elf transcription factors to the mixed cellular state seen upon CAF-1 suppression (Figure. 3). Finally, we also looked for motifs overrepresented in Hoxa9 OFF-specific ATAC-seq peaks and found as the most highly enriched sequence a consensus binding motif of the interferon regulatory factor 1, IRF1 (Figure. 4f). Several reports document a role for IRF1 in granulocyte differentiation(Abdollahi et al., 1991, Langlais et al., 2016), suggesting that IRF1 might contribute to commitment of GMPs to neutrophils upon Hoxa9 inactivation.

These analyses identify distinct transcription factors as candidate critical regulators of the observed cellular fates: C/EBPα as a driver during the initial phases of GMPs differentiation in both conditions, ELF family members as contributors to the establishment of the mixed cellular state upon suppression of CAF-1, and IRF1 as a Hoxa9-controlled factor that promotes terminal cell differentiation to neutrophils.

### C/EBPα and ELF1 transcription factors are required for the mixed cellular state triggered by CAF-1 suppression

To directly investigate a potential requirement of C/EBPα and ELF1 for the mixed cellular state upon depletion of CAF-1, we performed loss-of-function studies in GMPs using RNAi or CRISPR/Cas9-assisted gene editing (Figure. 5a-b). Consistent with C/EBPα motif enrichment in the commonly gained ATAC-seq sites, its depletion substantially impaired granulocyte differentiation initiated by loss of either CAF-1 or Hoxa9 (Figure. 5c-e and Figure5-figure supplement 1a-c). Upon C/EBPα knockdown we detected up to 36% or 16% fewer cells expressing Lys-GFP after 48h of treatment with IPTG or estrogen withdrawal, respectively (Figure. 5e). In contrast, since ELF1 motif is uniquely enriched in the CAF-1 OFF gained ATAC-seq sites its loss affected granulocytes marker activation upon CAF-1 suppression only (Figure. 5f-h and Figure5-figure supplement 1d-f). Upon ELF1 knockdown we detected up to 29% reduction in Lys-GFP positive cells at 48hrs, while having no observable effect upon HOXA9 inactivation (Figure. 5h). These results validate the above speculation and suggest that C/EBPα operates during the onset of differentiation in CAF-1 OFF and Hoxa9 OFF cells, whereas ELF1 specifically promotes differentiation initiated by CAF-1 depletion.

**Figure 5:**
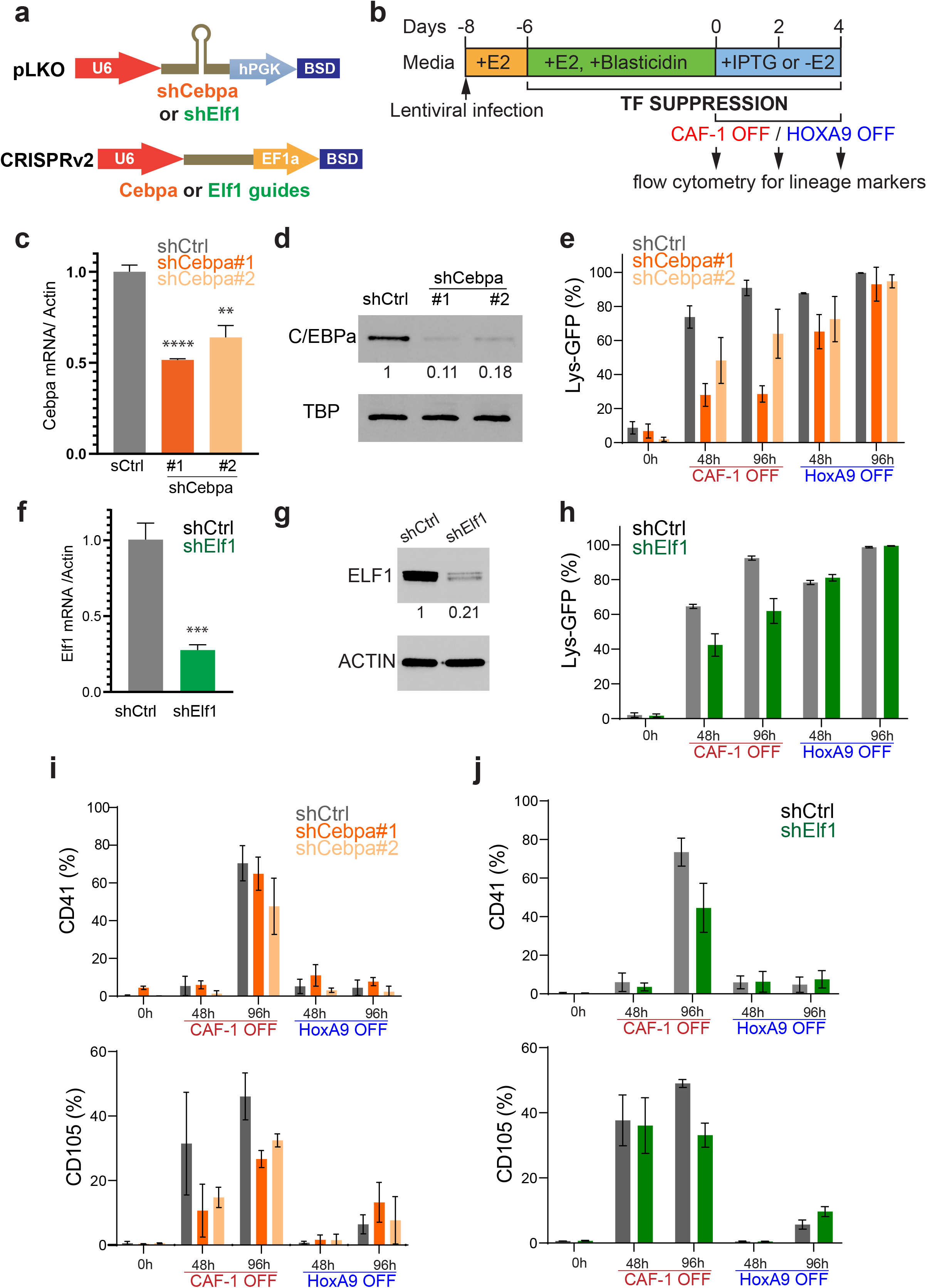
C/EBPα and ELF1 transcription factors are required for the mixed cellular state triggered by CAF-1 suppression. **a,** Schematic of Cebpa or Elf1 shRNAs cloned in pLKO.1 vector (top) and Cebpa or Elf1guides cloned in LentiCRISPR v2 vector. **b,** Strategy for loss of function analysis of C/EBPα and ELF1 in the context of CAF-1 OFF or HOXA9 OFF GMPs followed by assessment of lineage markers by flow cytometry. **c,d,f,g** confirmation of C/EBPα or ELF1 shRNA mediated suppression by quantitative RT-PCR and western blot analyses. **e&h**, Time course flow cytometric analysis of Lys-GFP expression in CAF-1 OFF versus HOXA9 OFF GMPs upon C/EBPα (orange,**e**) or ELF1 (green,**h**) knockdown as measure of granulocytes differentiation. **i-j,** Time course flow cytometric analysis of CD41 (top graphs) and CD105 (bottom graphs) expression in CAF-1 OFF versus HOXA9 OFF GMPs upon C/EBPα (**i**) or ELF1 (**j**) knockdown as measure the mixed cellular state observed in the CAF-1 OFF only condition.

To test the hypothesis that C/EBPα and Elf1 might contribute to the mixed cellular state of CAF-1 depleted cells, we monitored the effect of their individual suppression on induction of megakaryocyte (CD41) and erythrocyte (CD105) markers. Knockdown of either C/EBPα or Elf1 in CAF-1 OFF but not in Hoxa9 OFF cells led to the suppression of both markers, particularly at the 96hrs time point (Figure. 5i-j). For example, at 96hrs ELF1 loss in CAF-1 OFF cells resulted in 27% and 16% reduction of CD41 and CD105 positive cells, respectively (Figure. 5j). We conclude that suppression of CAF-1 overcomes the maturation block in GMPs, at least in part through facilitating access to C/EBPα and Elf1 transcription factors to their target sites. This, in turn, results in aberrant myeloid differentiation and a mixed cellular state (Figure5-figure supplement 2).

## Discussion

Mechanisms that sustain GMP identity are poorly understood. We discovered a role of restricted chromatin accessibility in safeguarding GMPs from a mixed cellular state by 1) dissecting the early and late transcriptional changes at a single cell level, 2) mapping accessible binding motifs of transcription factors upon CAF-1 suppression in comparison to canonical differentiation, and 3) interrogating the function of the identified candidate transcription factors. By combined analysis of single cell transcriptomes and chromatin accessibility manipulation by targeting CAF-1, we identified C/EBPα and ELF1 as key regulators in myeloid lineage commitment. It is worth noting that expression profiles of these transcription factors do not stand out in the analysis of transcriptomes alone. In fact, during the normal process of differentiation following inactivation of Hoxa9, C/EBPα expression levels decrease, while the levels of both C/EBPα and ELF transcription factors remain unchanged throughout the differentiation process upon CAF-1 repression (Figure4-figure supplement 2a-b). Thus, our analysis of chromatin accessibility in the context of modulating CAF-1 expression was instrumental to sense the activities of the critical transcription factors during myeloid differentiation.

scRNA-seq has deepened our understanding of transcriptional programs that govern lineage commitment and unmasked the heterogeneity even within sorted populations of cells. Consistent with our findings, a study employing scRNA-seq demonstrated that mixed lineage states do not arise during normal differentiation of myeloid progenitors (Paul et al., 2016). Additionally, this same study identified C/EBPa as a key regulator in lineage restriction, consistent with our identified role of C/EBPa in promoting differentiation of GMPs upon CAF-1 suppression or Hoxa9 inactivation. Based on these results and previous studies of CAF-1 dependent regulation of cellular plasticity(Cheloufi et al., 2015, Ishiuchi et al., 2015, Cheloufi and Hochedlinger, 2017, Ng et al., 2019), we speculate that perturbing CAF-1 may present a broadly applicable approach to discern transcriptional programs that are central to differentiation of various precursor cells into progressively more lineage-restricted progeny. This is noteworthy since most previous studies that investigated the function of CAF-1 in maintenance of cell identity did not probe the effect of CAF-1 ablation in progenitor – differentiated cell paradigms (Ishiuchi et al., 2015, Cheng et al., 2019, Ng et al., 2019, Volk et al., 2018).

Given that CAF-1 is essential for cell growth, studying its function in specific blood populations in-vivo during the course of differentiation remains challenging. The conditional knockout of CAF-1 within the hematopoietic compartment results in pancytopenia and the loss of hematopoietic progenitors in the bone marrow(Volk et al., 2018). Our study tests the direct role of CAF-1 in GMPs specifically without perturbing other stem and progenitor precursors. Importantly, the Hoxa9-GMPs captured in culture are karyotypically normal and retain the canonical myeloid differentiation potential observed in normal human and mouse hematopoesis(Sykes et al., 2016).

CAF-1 restricts chromatin accessibility both through its nucleosome assembly function and through recruitment of chromatin modifying enzymes that promote gene silencing. CHAF1b was recently proposed to compete against transcription factor binding, including C/EBPa in an MLL/AF9 leukemia system without affecting chromatin accessibility(Volk et al., 2018). However, in this study CHAF1B binding to chromatin was interrogated in a setting where Chaf1b overexpression alone was sufficient to enhance leukemogenesis. Here we focus on the gain in global chromatin accessibility sites upon CAF-1 knockdown in a normal myeloid progenitor state to predict which transcription factors become active and could therefore play a role in lineage plasticity. C/EBPα appears to be a common driver of cellular differentiation in both the setting of CAF-1 knockdown as well as Hoxa9 inactivation. However, in the context of CAF-1 suppression alone, we found that C/EBPα together with additional activation of other transcription factors such as ELF1 contribute to aberrant differentiation, resulting in a mixed cellular state. We speculate that CAF-1 restricts lineage choice in normal homeostasis through its nucleosome assembly function by blocking the binding sites of transcription factors that promote alternate lineages. Further studies are needed to understand how transcription factor binding and activity is modulated upon manipulation of CAF-1 in normal homeostasis and disease settings.

## Materials and Methods

### Cell culture and media

Granulocyte-macrophage progenitor cells containing *Hoxa9::ER* were cultured in RPMI-1640 supplemented with 10% FBS, 100U ml^−1^ penicillin, 100 μg ml^−1^ streptomycin, 2mM L-alanyl-L-glutamine dipeptide and stem cell factor (SCF). The source of SCF was conditioned media generated from a Chinese hamster ovary (CHO) cell line that stably secretes SCF. Conditioned medium was added at a final concentration of 1 - 2 % depending on the batch. β-Estradiol (abbreviated E2) was added to a final concentration of 150 ng/ml from a 3 mg/ml stock dissolved in 100% ethanol. Packaging cells (293T cell line) for producing lentiviral particles were cultured in DMEM supplemented with 10% FBS, 100U ml^−1^ penicillin, 100 μg ml^−1^ streptomycin, 55 μM β-mercaptoethanol, 2 mM L-alanyl-L-glutamine dipeptide and 1x MEM NEAA. All cells were cultured at 37°C with 5% CO_2_. Cells were tested for mycoplasma contamination and found to be negative.

### Flow cytometry

Antibodies (mouse: CD11b, Gr-1, CD41, CD105) were all purchased from BioLegend. Cells were suspended in FACS buffer (PBS + 5% FBS + 1 mM EDTA) and stained for 20 min at 4°C in the dark. Propidium iodide was used as viability dyes to help identify dead cells prior to flow cytometry. Flow cytometry assays at UCR were performed on NovoCyte flow cytometer, acquired using NovoExpress software, and analyzed using FlowJo software. Flow cytometry assays at MGH were performed on the BD LSR2 flow cytometer, acquired using Diva software, and analyzed using FlowJo software.

### Time course, point of commitment and growth curve

For IPTG-inducible Chaf1b shRNA expression (CAF-1 OFF), IPTG was added at a concentration of 500 μM. For Hoxa9 inactivation, cells were washed with PBS twice and then cultured in media without E2. Cell media were replenished with corresponding media every two days during time courses. In point of commitment experiment, cells were first cultured with IPTG addition or E2 withdrawal with 24hrs pulse increments, followed by IPTG withdrawal or E2 addition to deactivate Chaf1b shRNA or activate Hoxa9. Growth curves were performed using ViaLight Plus Cell Proliferation and Cytotoxicity BioAssay Kit (Lonza) according to the supplier’s instruction.

### Quantitative RT-PCR

RNA was extracted with the Direct-zol RNA miniprep plus kit (Zymo Research) and then reverse transcribed with the PrimeScript RT reaction kit (TaKaRa) according to the supplier’s instruction. Quantitative PCR was performed using the PowerUp SYBR Green master mix (Applied Biosystems) in a BIO-RAD CFX connect cycler. Primers used were: Cebpa-F: ATAGACATCAGCGCCTACATCGA; Cebpa-R: GTCGGCTGTGCTGGAAGAG; Elf1-F: TGCAAGTAACGGCATGGAGG; Elf1-R: AGGAACATGTTCCACAATAACAGCA; Pf4-F: CCGAAGAAAGCGATGGAGATCT; Pf4-R: ATTCTTCAGGGTGGCTATGAGC; Cd41-F: TGCCGTGGTATTGCATGGA; Cd41-R: CAGACAAGCCTCTCAAAGCCCT; Cd105-F: GTACCCACAAGTCTCGCAGAA; Cd105-R: AGATGTGACAGCATTCCGGG; β-actin-F: CGCCACCAGTTCGCCATGGA; β-actin-R: TACAGCCCGGGGAGCATCGT. Results are presented as 2^−ΔΔCt^ values normalized to the expression of β-actin and negative control samples. All reactions were performed in triplicate. The means and standard deviations were calculated in GraphPad Prism 8 software.

### SDS-PAGE and western blot analysis

Whole-cell lysates were run on 10 or 15% SDS-polyacrylamide gels and transferred to nitrocellulose membrane (1620097, Bio-Rad) by standard methods. Membranes were blocked for 1h for 5% non-fat dry milk in 1 ×TBS with 0.05% Tween-20 (TBST), rinsed, and incubated with primary antibody diluted in 3% BSA in TBST overnight at 4 °C. The following primary antibodies were used: anti-Chaf1a (sc-10206, Santa Cruz, discontinued), anti-Chaf1b (sc-393662, Santa Cruz), anti-TBP (ab818, Abcam), anti-CEBPα (8178, Cell Signaling), anti-ELF1(sc-133096, Santa Cruz). Blots were washed in TBST, incubated with HRP-conjugated secondary antibodies for semi-quantitative western blot analysis. Secondary antibodies were incubated in 5% milk in TBST for 1 h at room temperature and washed again. HRP signal was detected by Western Lightning Plus-ECL (NEL103E001EA, Perkin Elmer).

### Cytospins and Wright-Giemsa Staining

Cells were prepared in PBS at a concentration of approximately 2 million/ml. Cytospin (Thermo Scientific Shandon) preparations were made (1,000 rpm, 60 s), and the cells were allowed to air dry. Cells were stained in 100% Wright-Giemsa (Siemens) for 2 min, and in 20% Wright-Giemsa diluted in buffer for 12 min. Stained cells were rinsed in deionized water, and coverslips were affixed with Permount prior to microscopy.

### Phagocytosis Assay

ER-Hoxa9 cells were differentiated out of E2 for a period of 4 days. Cells were resuspended in media along with fluorescein-labeled heat-killed Escherichia coli BioParticles (pHrodo, Molecular Probes). Cells and bioparticles were agitated at 37°C for 60 min prior to flow cytometry; DAPI was used as a viability dye.

### Lentivirus production

shRNAs targeting Chaf1b were cloned into pLKO-TRC025 vector (Broad institute) harboring a puromycin-selectable marker. shRNAs were used: shChaf1b#1 (TRCN0000318223): GCTGTCAATGTTGTACGCTTT; shChaf1b#2 (TRCN0000318224) CGTCATTCTGTTGTGGAAGAT. Chaf1b shRNA were shuttled into into pLKO-TRC912 1X LacI vector harboring a puromycin-selectable marker for inducible expression. shRNA targeting the C/EBPα and ELF1 were cloned into pLKO.1 vector (Addgene, plasmid # 26655) harboring a blasticidin-selectable marker. shRNAs were used: shCebpa#1: GCCGAGATAAAGCCAAACAAC; shCebpa#2: GGACAAGAACAGCAACGAGTA; shElf1: GTGATCCTGCTATATTTCCTG; shCTRL (scrambled control): CCTAAGGTTAAGTCGCCCTCGC. sgRNAs targeting the C/EBPα and ELF1 locus were cloned into lentiCRISPR v2 vector (Addgene, plasmid #83480) harboring the wild type Cas9 coding region, an sgRNA expression cassette, and a blasticidin-selectable marker. Guide sequences used were: gCebpa#1: AGAAGTCGGCCGACTCCATG; gCebpa#2: GCGGCGCGGTCATGTCCGCG; gElf1: ATGAACAGTTCGGAAGAGCT. Lentiviruses were produced by transfection of 293T cells with Δ8.9 and VSVG plasmids. Virus was harvested at 36, 60 and 84h post-transfection and precipitated using PEG3500 (Sigma-Aldrich, cat#P4338). For transduction, 150,000 cells were plated per well in a 12-well dish, and spin infected at 2500 rpm for 1.5h in the presence of 10 μg mL^−1^ of polybrene (Millipore). After 48h, transduced cells were selected with 10μg mL^−1^ blasticidin (Gibco) for 6 days.

### 10x Chromium single-cell RNA Sequencing

Hoxa9-GMP cells were collected at 0h, 48hrs and 96hrs upon CAF-1 suppression or Hoxa9 inactivation. Cells were then washed and resuspended with 10% FBS. The subsequent preparation was conducted following the instruction manual of 10X Chromium v2. The cDNA library and final library after index preparation were checked with bioanalyzer (High Sensitivity DNA reagents, Agilent Technology) for quality control. Following the library preparation, the sequencing was performed with paired-end sequencing of 75nt each end on HiSeq4000, by Novo Gene, Inc. Cells were sequenced to an average depth of 100,000 reads per cell.

### 10X Chromium single-cell RNA Seq bioinformatics analysis

The raw reads were mapped onto the mouse genome Ensembl gene model file Mus_musculus.GRCm38.gff using a standard CellRanger 3.1.0 pipeline. The package Seurat v3.0.1 was used in this study. Cells with less than 200 genes or more than 6000 genes (potential doublet) were removed, and only genes that were found expressed in at least 3 cells were kept in the analysis. Cells with mitochondrial gene percentage >10% were filtered out. Unsupervised clustering analysis of 10X scRNAseq data performed after normalizing for total reads per cell and log transformed.

### Bulk RNA-seq

Total RNA from cells was DNAse treated and purified using Qiagen RNeasy mini kit (according to manufacturer’s instructions). RNA quality was assessed using an Agilent 2100 Bioanalyzer. RNA-seq libraries were prepared with NEBNext UltraDirectional kit (New England Biolabs). Libraries were pooled and sequenced on Illumina HiSeq 2500 instrument. On average 28 million reads were generated per library.

### Bioinformatics analysis of bulk RNA-seq data

STAR aligner(Dobin et al., 2013) was used to map sequencing reads to transcripts in the mouse mm9 reference genome. Read counts for individual transcripts were produced with HTSeq-count(Anders et al., 2015), followed by the estimation of expression values and detection of differentially expressed transcripts using EdgeR(Robinson et al., 2010). Differentially expressed genes were defined by at least 2-fold change with FDR less than 0.01. GSEA(Subramanian et al., 2005) and EnrichR was used to analyze the enrichment of functional gene groups among differential expressed genes.

### Bulk ATAC-seq

To generate ATAC-seq libraries, 50,000 cells were used and libraries were constructed as previously described(Buenrostro et al., 2013). Briefly, cells were washed in PBS twice, counted and nuclei were isolated from 100,000 cells using 100 μl hypotonic buffer (10 mM Tris pH 7.4, 10 mM NaCl, 3 mM MgCl_2_, 0.1% NP40) to generate two independent transposition reactions. Nuclei were split in half and treated with 2.5 μl Tn5 Transposase (Illumina) for 30 min at 37 °C. DNA from transposed nuclei were then isolated and PCR-amplified using barcoded Nextera primers (Illumina). Library quality control and quantitation was carried out using high-sensitivity DNA Bioanalyzer and Qubit, followed by paired-end sequencing (PE50) on the Illumina HiSeq 2500 instrument.

### Bioinformatics analysis of bulk ATAC-seq data

Sequencing reads were aligned against the mm9 reference genome using BWA(Li and Durbin, 2010). Reads mapping to the mitochondrial genome and duplicate reads were removed. Peaks were called using the HOTSPOT method(John et al., 2011). Read numbers and densities were calculated across all samples for each genomic region among the union of all peaks in all samples. Regions of differential accessibility were identified using edgeR(Robinson et al., 2010), with the cutoffs of at least 2-fold difference and FDR<0.01. Sequence motifs enriched among these differentially accessible regions were identified using MEME-ChIP(Bailey et al., 2009).

### Statistical analysis

Unless otherwise specified, values are depicted as mean ± SD. Parameters including statistical significance, P-value and statistical analysis methods are reported in the Figure legends and Supplementary Figure legends. Statistical analysis was performed using Student’s t-test. In all cases, \**P* <0.05, \*\**P* < 0.01, \*\*\**P* < 0.001, \*\*\*\**P* < 0.0001 were considered statistically significant.

## Acknowledgments

We thank Xiwei Wu at COH for support with 10X genomics library construction, Novogene for single cell RNA-seq, Rachel Behar at the UCR stem cell core, Marisol Arellano and David Lo at the UCR school of medicine for support with flow cytometry, Glenn Cowley and David Root at the Broad Institute for support with inducible shRNAs expression, the Hochedlinger, Murn, Martinez, Chen and Karginov labs for discussions, members of the Cheloufi and Murn labs for critical reading of the manuscript, the MGH CRM/HSCI Flow Core; and the MGH Molecular biology department genomics core facility. We are grateful to Dr. R. Luben and F. Sladek and A. Ray for guidance. SC was supported by PRCRP at the Department of Defense (CA 120212) and MGH ECOR fellowships, SC and KH were supported by the NIH (5R01HD058013-10). SC & YG are supported by the City of Hope/UCR biomedical research initiative (CUBRI) and UC cancer research coordinating committee (CRCC) seed grants.

## Author contributions

SC, DBS and KH designed the study. SC, DBS and YG performed and analyzed the experiments. YG, FJ, RS, DF, MJ and RCR performed bioinformatics analysis. JM cloned shRNAs and Chaf1b overexpression constructs and helped with data interpretation. DBS provided the Hoxa9-GMP lines. CC helped with kinetics experiments. JM and MAB provided intellectual support. KH provided intellectual support and mentoring. SC, JM, DBS and YG wrote the manuscript with input from all co-authors.

## FIGURE SUPPLEMENTS

### Figure supplements legends

**Figure2-figure supplement 1:**
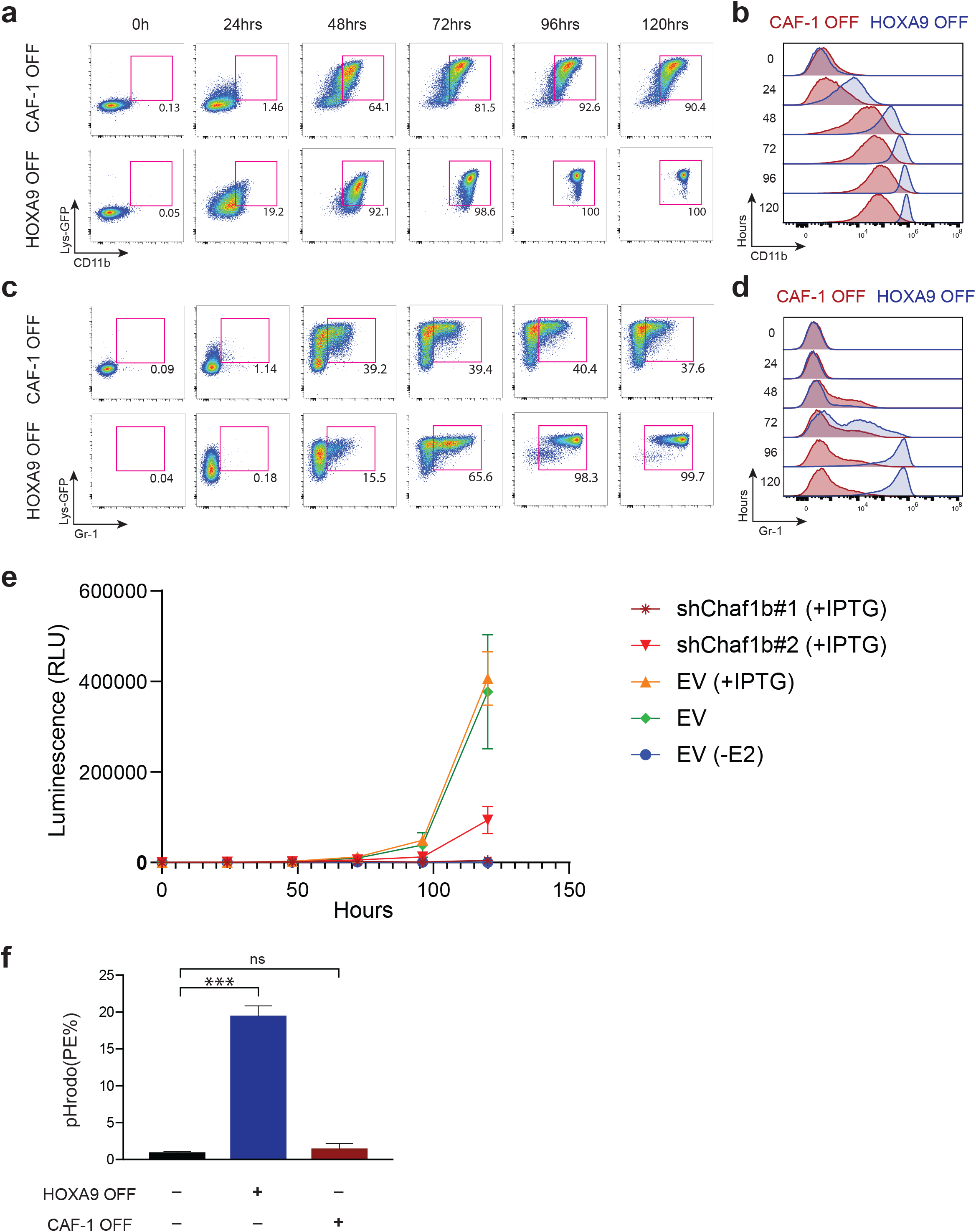
CAF-1 suppression mirrors canonical GMP differentiation but exhibit no phagocytotic activity. **a-d,** Representative flow cytometry plots and histograms of CD11b **(a, b)** and Gr-1 **(c, d)** expression during 5-day time course in CAF-1 OFF versus HOXA9 OFF GMPs. **e,** Growth curves comparing HOXA9 OFF versus CAF-1 OFF GMPs in two IPTG inducible sub-clones expressing Chaf1b shRNA. **f**, Quantification of phagocytotic activities induced HOXA9 OFF versus CAF-1 OFF GMPs measured from the flow cytometric assay shown in **Figure. 2e.**

**Figure3-figure supplement 1:**
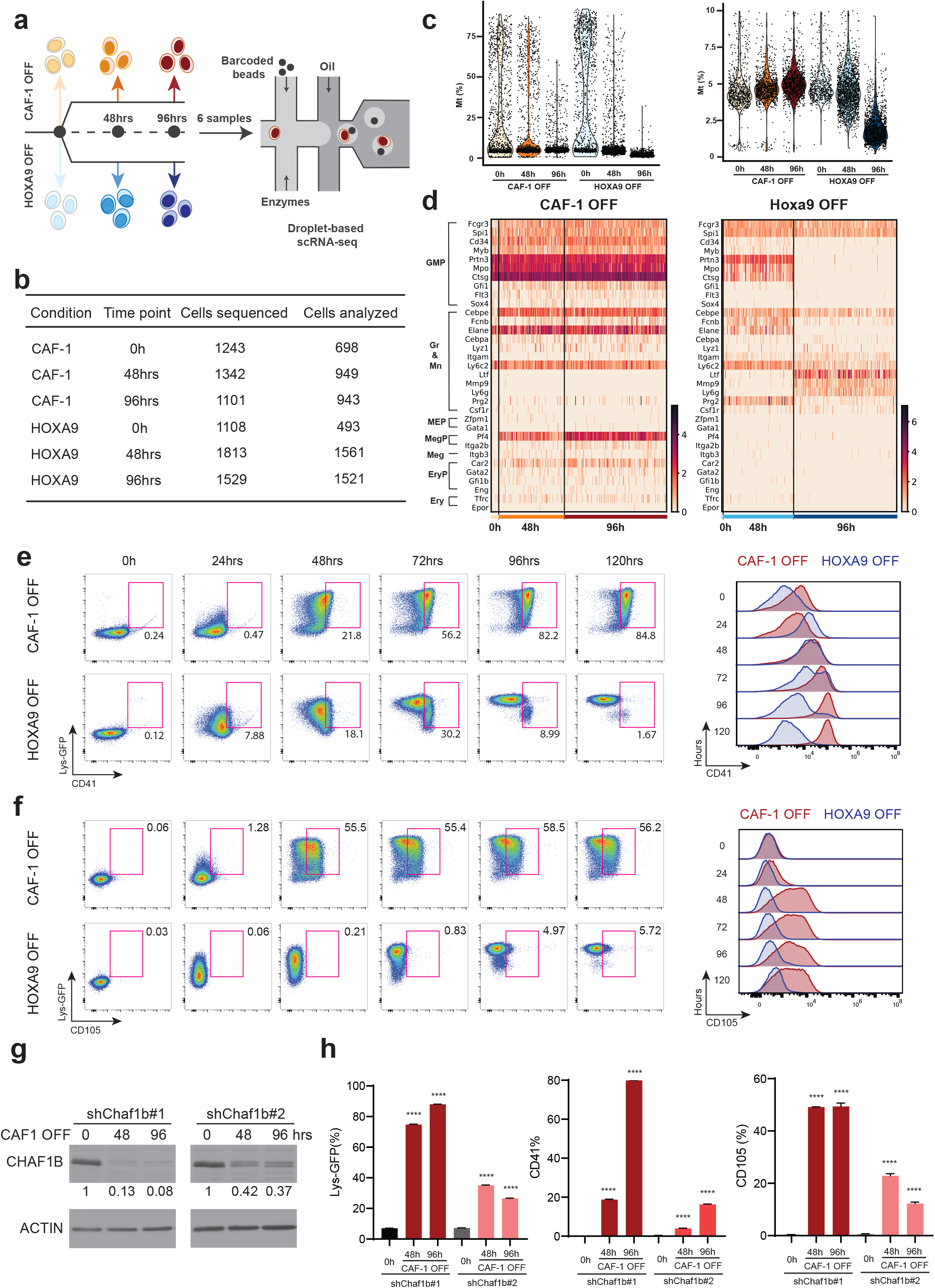
Activation of mixed lineage genes upon CAF-1 suppression in GMPs. **a,** Experimental design for scRNA-seq experiments depicting the time course analysis at 48 and 96 hours. **b**, Summary of number of single cells sequenced and analyzed in each condition and time point. **c,** Percentage of mitochondrial UMIs per cell per sample before and after filtering. **d,** Heatmap displaying gene expression patterns of a curated set of lineage specific genes (surface markers and transcription factors) at different time points in CAF-1 OFF versus HOXA9 OFF cells genes shown in **Figure. 3c&d** only in Gr-1 positive cells confirming the mixed cellular state in CAF-1 OFF only condition. **e, f** Flow cytometric plots and histograms of Lys-GFP, CD41 and CD105 expression showing a more detailed time course over 5days in CAF-1 OFF versus HOXA9 OFF cells reflecting the mixed cellular state of CAF-1 OFF cells only. Lys-GFP and CD41 or CD105 double positive cells are gated in the dot plots and the expression levels of the aberrantly expressed lineage markers CD41 and CD105 are represented in the histogram. **g,** Western blot analysis showing different levels of CHAF1B reduction in a strong versus weak IPTG inducible sub-clone (see corresponding growth curve in Supplemental Fig. S1e). **h** Flow cytometric analysis of Lys-GFP, CD41 and CD105 of the CAF-1 OFF independent sub-clones in **g** showing a dosage dependent activation of the aberrantly expressed lineage markers upon CAF-1 suppression.

**Figure3-figure supplement 2:**
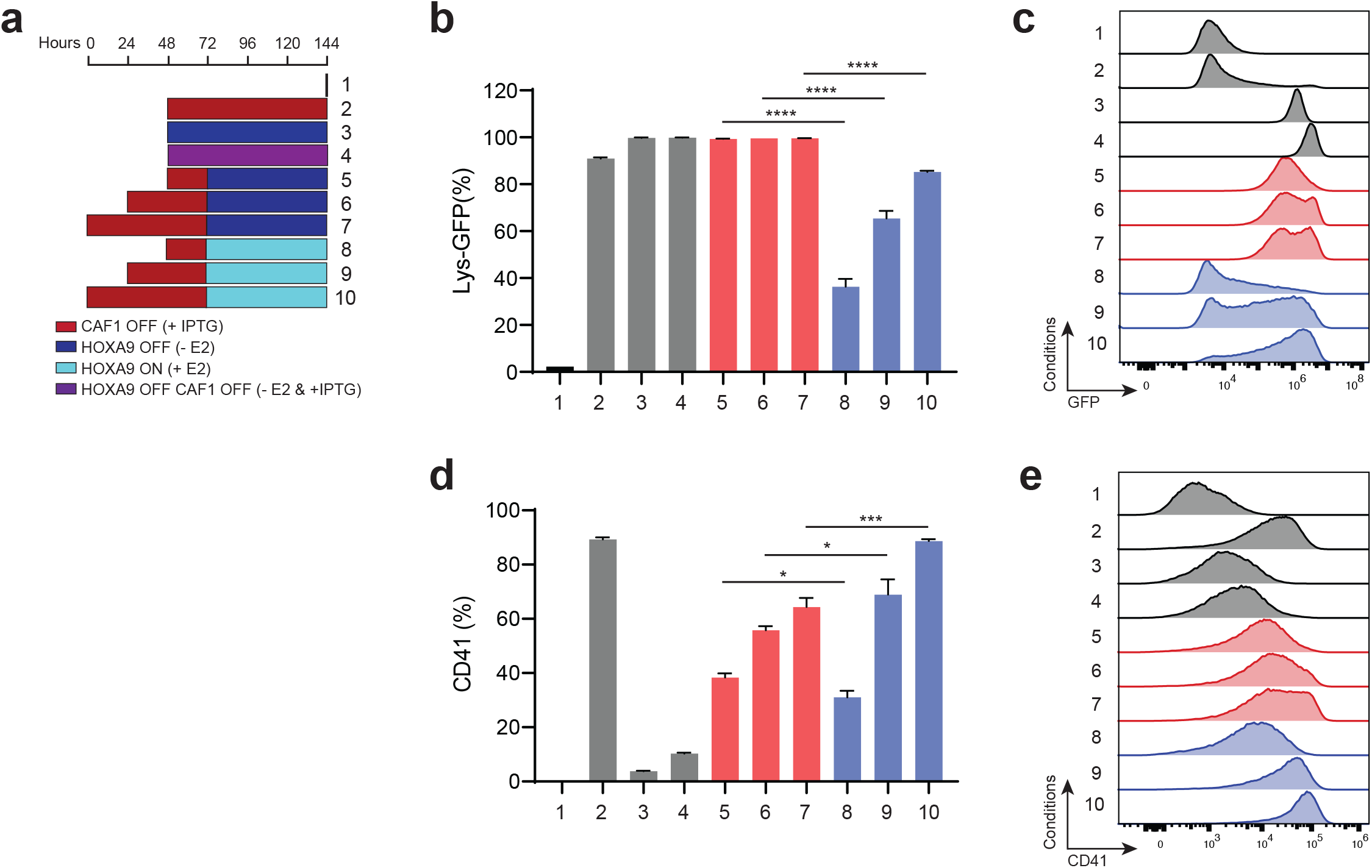
The mixed cellular state upon CAF-1 suppression is not reversed with HOXA9 inactivation. **a,** Pulse strategy showing sequential inactivation of CAF-1 and HOXA9. Individual IPTG and -E2 and simultaneous pulses are performed as controls. **b**, Flow cytometric analysis of Lys-GFP positive cells in all pulses shown in **a**. **c**, Representative flow cytometry histograms showing intensity Lys-GFP expression for all corresponding pulses. **d,** Flow cytometric analysis of CD41 positive cells of all pulses shown in (**a**) as a readout of mixed cellular state upon CAF-1 suppression. **e**, Representative flow cytometry histograms showing intensity of CD41 expression for all corresponding pulses.

**Figure4-figure supplement 1:**
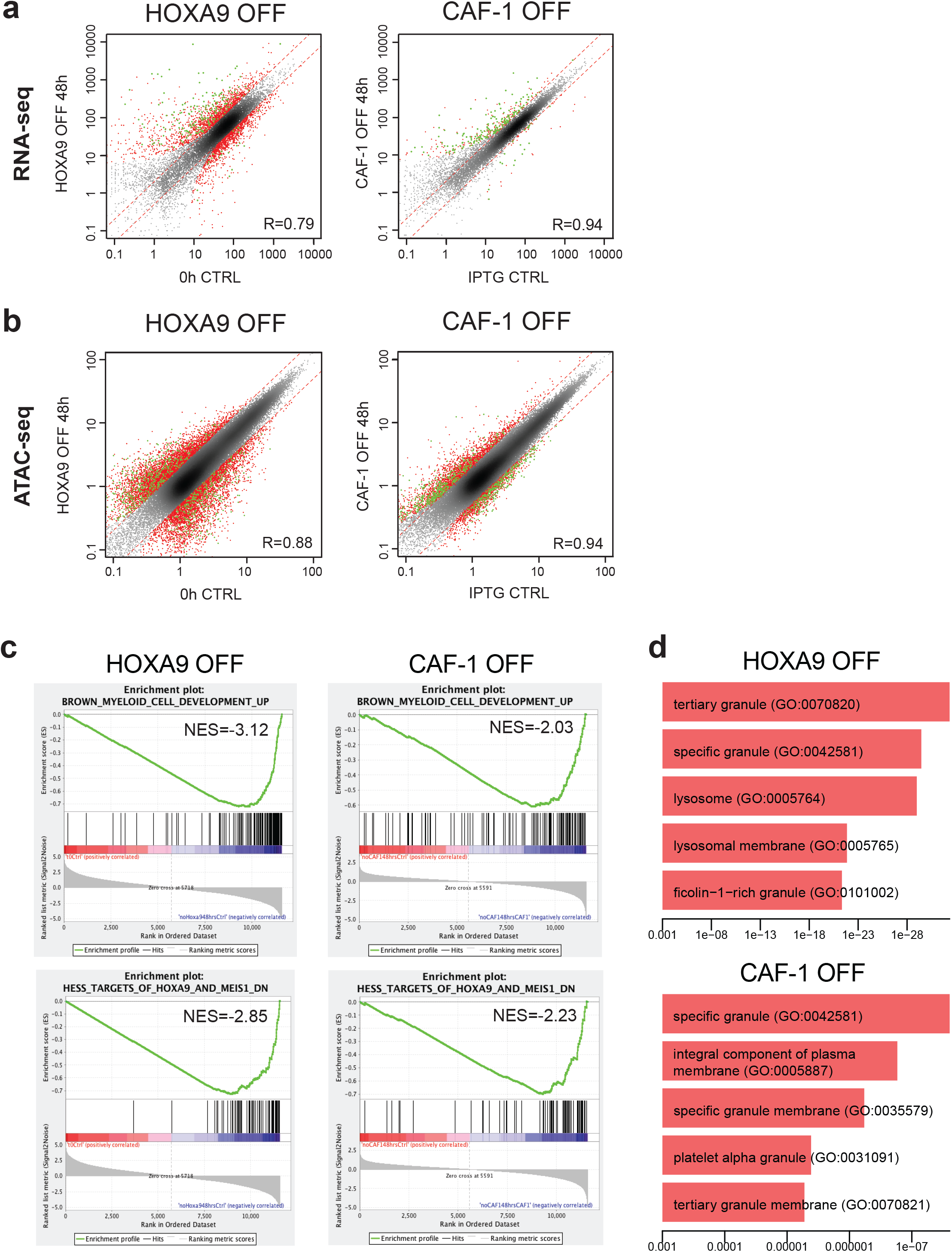
Magnitude of gene expression and chromatin accessibility changes and gene set enrichment analysis comparing CAF-1 OFF and HOXA9 OFF GMPs. **a&b** Gene expression levels and ATAC-seq peak intensity comparing untreated (0h, +E2 control or IPTG CTRL) versus inactivation of either HOXA9 or CAF-1 at 48hrs. Genes or peaks off the diagonal dotted lines are differentially expressed. Uniquely regulated genes or peaks in each condition are shown in red and commonly regulated gene or peaks are shown in green (compare to venn diagrams in Fig.4 a&b). **c**, GSEA of bulk RNA-seq data at 48hrs in CAF-1 OFF versus HOXA9 OFF GMPs showing similar significant sets recovered in both conditions reflecting myeloid differentiation. **d,** EnrichR GO cellular component of bulk RNA-seq data at 48hrs in CAF-1 OFF versus HOXA9 OFF GMPs depicting significant enrichment of gene sets associated with granulocytes differentiation.

**Figure4-figure supplement 2:**
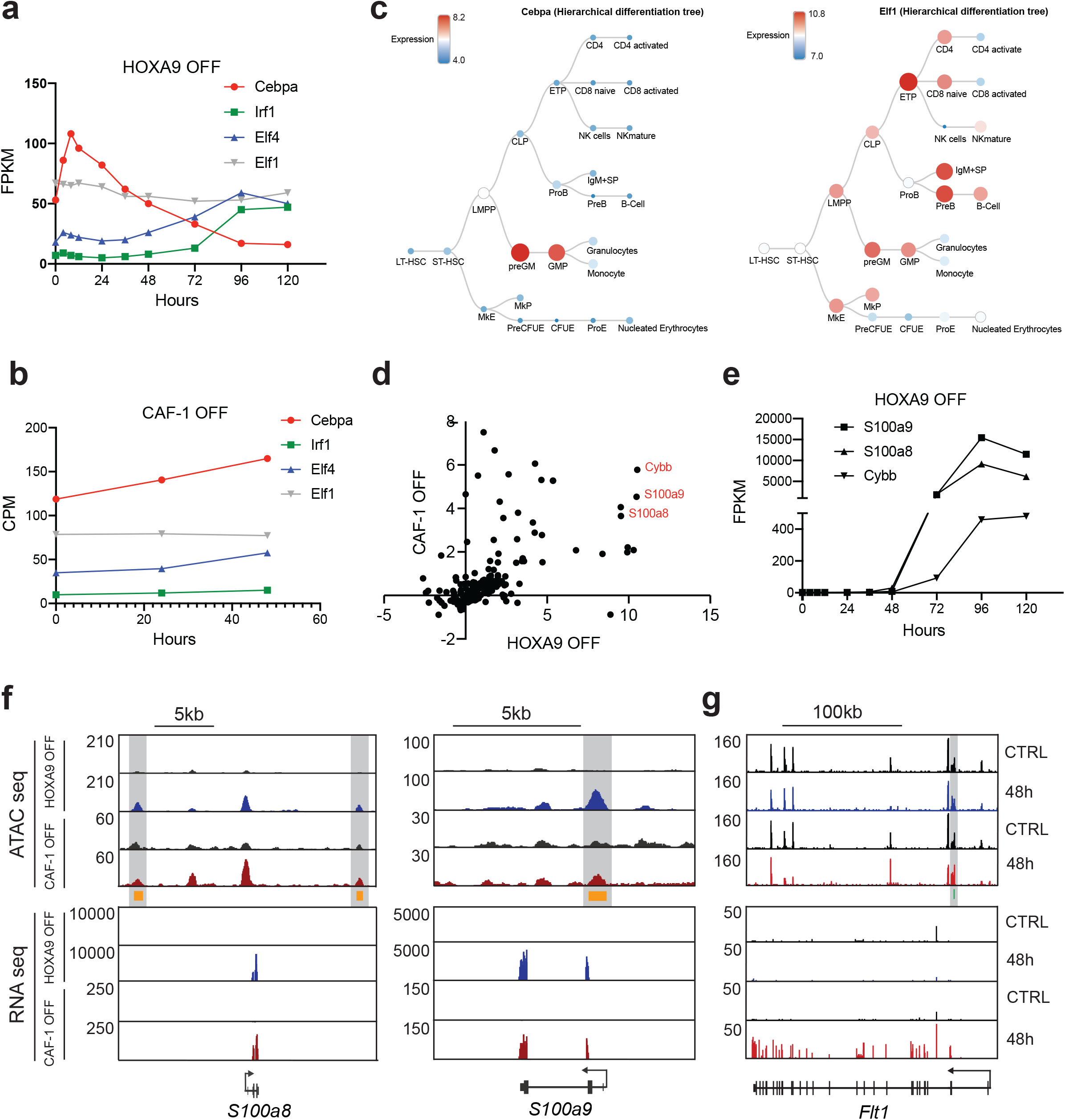
Chromatin accessibility changes upon CAF-1 perturbation reveals common and alternate transcriptional programs. **a,** Normalized fragments per kilobase of transcript per million mapped reads (FPKM) values of transcription factors Cebpa, Irf1, Elf4 and Elf1 during Hoxa9 inactivation from previously published canonical GMP differentiation dataset(Sykes et al. 2016). **b,** Count per million reads (CPM) of transcription factors Cebpa, Irf1, Elf4 and Elf1 at 0h, 24hrs and 48hrs upon CAF-1 suppression showing static expression of candidate transcription factors. **c,** Cebpa and Elf1 expression profile in hematopoietic hierarchical trees from blood spot database(Bagger et al. 2016). **d,** Correlation analysis of differentially expressed genes at 48hrs that are within 100KB distance from the predicted C/EBPα gained ATAC-seq sites in CAF-1 OFF versus HOXA9 OFF conditions. **e,** FPKM values of top commonly regulated targets genes (S100a9, S100a8 and Cybb) neighboring C/EBPα motif ATAC-seq peaks over time from previously published canonical GMP differentiation dataset(Sykes et al. 2016). **f&g** Representative ATAC-seq and RNA-seq genome snap shots depicting the correlation between the gain in chromatin accessibility and activation of neighboring genes. **f,** examples of commonly targeted genes between HOXA9 OFF and CAF-1 (S100a9, S100a8) neighboring CEBPα (orange) predicted binding sites. **g**, example of a CAF-1 OFF uniquely targeted gene Flt1 neighboring ELF1 predicted binding site (green).

**Figure5-figure supplement 1:**
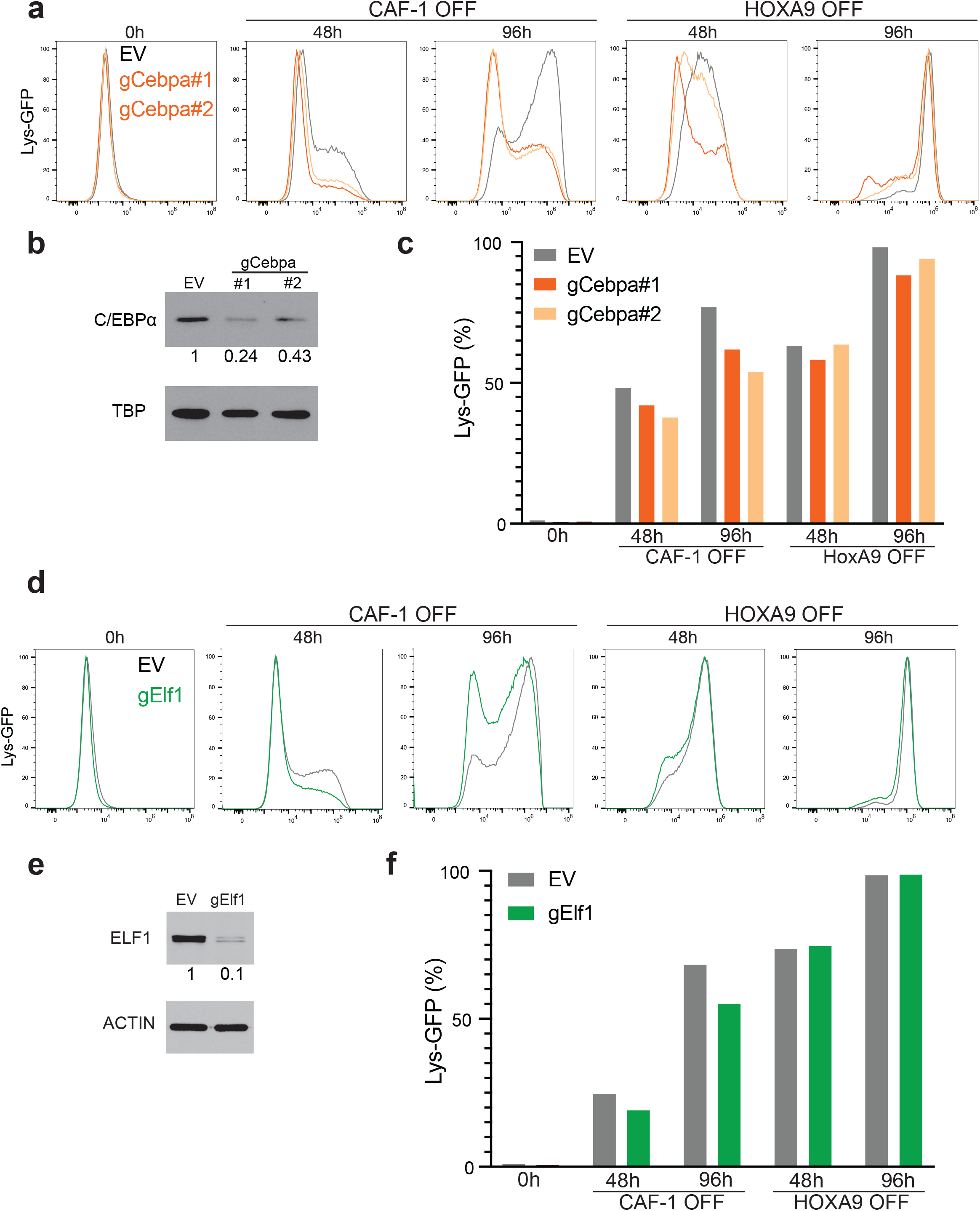
CRISPR/Cas9-assisted gene editing of C/EBPα and ELF1 transcription factors in HOXA9 OFF and CAF-1 OFF GMPs. **a,** Flow Cytometry histograms reflecting Lys-GFP expression changes upon C/EBPα deletion at 0h, 48hrs and 96hrs in CAF-1 OFF versus HOXA9 GMPs. **b,** Western blot analysis confirming reduced C/EBPα protein levels mediated by two independent guide RNAs. **c,** Quantification of Lys-GFP positive cells shown in **a**. **d,** Flow Cytometry histograms reflecting Lys-GFP expression changes upon ELF1 deletion at 0h, 48hrs and 96hrs in CAF-1 OFF versus HOXA9 GMPs. **e,** Western blot analysis confirming reduced ELF1 at protein level mediated by Elf1 guide RNA. **f**, Quantification of Lys-GFP positive cells shown in **d**.

**Figure5-figure supplement 2:**
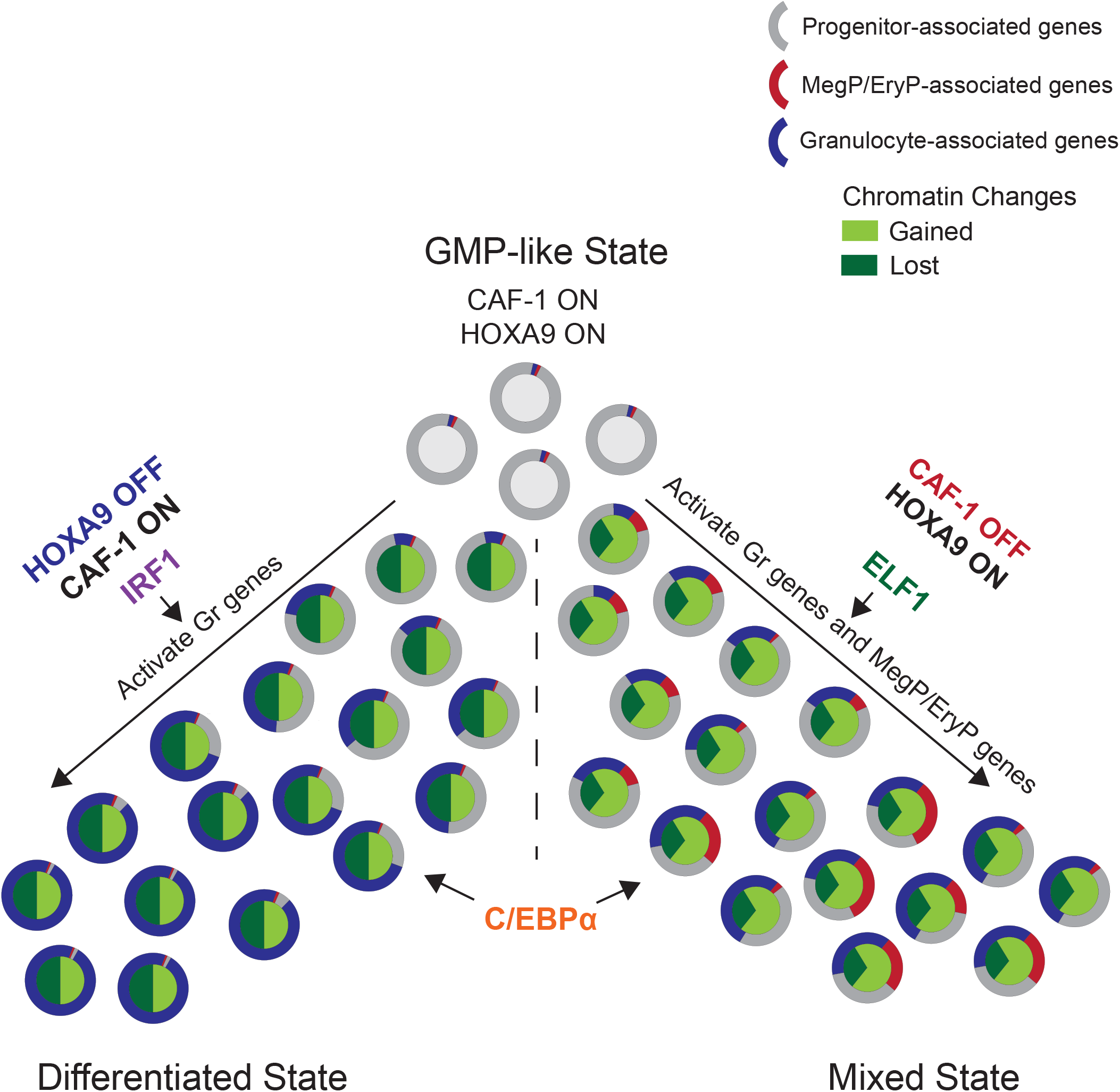
Proposed model of how chromatin accessibility manipulation affects normal and aberrant GMPs differentiation. Left branch depicts GMPs canonical differentiation upon Hoxa9 inactivation and the action of C/EBPα and IRF1 transcription factors. Right branch depicts aberrant differentiation upon CAF-1 suppression and the action of C/EBPα and ELF1 resulting a mixed cellular state.

## References

Abdollahi, A., Lord, K. A., Hoffman-Liebermann, B. & Liebermann, D. A. 1991. Interferon regulatory factor 1 is a myeloid differentiation primary response gene induced by interleukin 6 and leukemia inhibitory factor: role in growth inhibition. Cell Growth Differ, 2, 401–7.

Anders, S., Pyl, P. T. & Huber, W. 2015. HTSeq--a Python framework to work with high-throughput sequencing data. Bioinformatics, 31, 166–9.

Atlasi, Y. & Stunnenberg, H. G. 2017. The interplay of epigenetic marks during stem cell differentiation and development. Nat Rev Genet, 18, 643–658.

Bailey, T. L., Boden, M., Buske, F. A., Frith, M., Grant, C. E., Clementi, L., Ren, J., Li, W. W. & Noble, W. S. 2009. MEME SUITE: tools for motif discovery and searching. Nucleic Acids Res, 37, W202–8.

Becht, E., Mcinnes, L., Healy, J., Dutertre, C. A., Kwok, I. W. H., Ng, L. G., Ginhoux, F. & Newell, E. W. 2018. Dimensionality reduction for visualizing single-cell data using UMAP. Nat Biotechnol.

Buenrostro, J. D., Giresi, P. G., Zaba, L. C., Chang, H. Y. & Greenleaf, W. J. 2013. Transposition of native chromatin for fast and sensitive epigenomic profiling of open chromatin, DNA-binding proteins and nucleosome position. Nat Methods, 10, 1213–8.

Carrelha, J., Meng, Y., Kettyle, L. M., Luis, T. C., Norfo, R., Alcolea, V., Boukarabila, H., Grasso, F., Gambardella, A., Grover, A., Hogstrand, K., Lord, A. M., Sanjuan-PLA, A., Woll, P. S., Nerlov, C. & Jacobsen, S. E. W. 2018. Hierarchically related lineage-restricted fates of multipotent haematopoietic stem cells. Nature, 554, 106–111.

Cheloufi, S., Elling, U., Hopfgartner, B., Jung, Y. L., Murn, J., Ninova, M., Hubmann, M., Badeaux, A. I., Euong Ang, C., Tenen, D., Wesche, D. J., Abazova, N., Hogue, M., Tasdemir, N., Brumbaugh, J., Rathert, P., Jude, J., Ferrari, F., Blanco, A., Fellner, M., Wenzel, D., Zinner, M., Vidal, S. E., Bell, O., Stadtfeld, M., Chang, H. Y., Almouzni, G., Lowe, W., Rinn, J., Wernig, M., Aravin, A., Shi, Y., Park, P. J., Penninger, J. M., Zuber, J. & Hochedlinger, K. 2015. The histone chaperone CAF-1 safeguards somatic cell identity. Nature, 528, 218–24.

Cheloufi, S. & Hochedlinger, K. 2017. Emerging roles of the histone chaperone CAF-1 in cellular plasticity. Curr Opin Genet Dev, 46, 83–94.

Cheng, L., Zhang, X., Wang, Y., Gan, H., Xu, X., Lv, X., Hua, X., Que, J., Ordog, T. & Zhang, Z. 2019. Chromatin Assembly Factor 1 (CAF-1) facilitates the establishment of facultative heterochromatin during pluripotency exit. Nucleic Acids Res, 47, 11114–11131.

Dobin, A., Davis, C. A., Schlesinger, F., Drenkow, J., Zaleski, C., Jha, S., Batut, P., Chaisson, M. & Gingeras, T. R. 2013. STAR: ultrafast universal RNA-seq aligner. Bioinformatics, 29, 15–21.

Faust, N., Varas, F., Kelly, L. M., Heck, S. & Graf, T. 2000. Insertion of enhanced green fluorescent protein into the lysozyme gene creates mice with green fluorescent granulocytes and macrophages. Blood, 96, 719–26.

Gallant, S. & Gilkeson, G. 2006. ETS transcription factors and regulation of immunity. Arch Immunol Ther Exp (Warsz), 54, 149–63.

Hammond, C. M., Stromme, C. B., Huang, H., Patel, D. J. & Groth, A. 2017. Histone chaperone networks shaping chromatin function. Nat Rev Mol Cell Biol, 18, 141–158.

Heyworth, C., Pearson, S., May, G. & Enver, T. 2002. Transcription factor-mediated lineage switching reveals plasticity in primary committed progenitor cells. EMBO J, 21, 3770–81.

Houlard, M., Berlivet, S., Probst, A. V., Quivy, J. P., Hery, P., Almouzni, G. & Gerard, M. 2006. CAF-1 is essential for heterochromatin organization in pluripotent embryonic cells. PLoS Genet, 2, e181.

Ishiuchi, T., Enriquez-Gasca, R., Mizutani, E., Boskovic, A., Ziegler-Birling, C., Rodriguez-Terrones, D., Wakayama, T., Vaquerizas, J. M. & Torres-Padilla, M. E. 2015. Early embryonic-like cells are induced by downregulating replication-dependent chromatin assembly. Nat Struct Mol Biol, 22, 662–71.

Jacobsen, S. E. W. & Nerlov, C. 2019. Haematopoiesis in the era of advanced single-cell technologies. Nat Cell Biol, 21, 2–8.

John, S., Sabo, P. J., Thurman, R. E., Sung, M. H., Biddie, S. C., Johnson, T. A., Hager, G. L. & Stamatoyannopoulos, J. A. 2011. Chromatin accessibility pre-determines glucocorticoid receptor binding patterns. Nat Genet, 43, 264–8.

Kuleshov, M. V., Jones, M. R., Rouillard, A. D., Fernandez, N. F., Duan, Q., Wang, Z., Koplev, S., Jenkins, S. L., Jagodnik, K. M., Lachmann, A., Mcdermott, M. G., Monteiro, C. D., Gundersen, G. W. & Ma’ayan, A. 2016. Enrichr: a comprehensive gene set enrichment analysis web server 2016 update. Nucleic Acids Res, 44, W90–7.

Langlais, D., Barreiro, L. B. & Gros, P. 2016. The macrophage IRF8/IRF1 regulome is required for protection against infections and is associated with chronic inflammation. J Exp Med, 213, 585–603.

Laurenti, E. & Gottgens, B. 2018. From haematopoietic stem cells to complex differentiation landscapes. Nature, 553, 418–426.

Li, H. & Durbin, R. 2010. Fast and accurate long-read alignment with Burrows-Wheeler transform. Bioinformatics, 26, 589–95.

Liggett, L. A. & Sankaran, V. G. 2020. Unraveling Hematopoiesis through the Lens of Genomics. Cell, 182, 1384–1400.

Machanick, P. & Bailey, T. L. 2011. MEME-ChIP: motif analysis of large DNA datasets. Bioinformatics, 27, 1696–7.

Ng, C., Aichinger, M., Nguyen, T., Au, C., Najar, T., Wu, L., Mesa, K. R., Liao, W., Quivy, J. P., Hubert, B., Almouzni, G., Zuber, J. & Littman, D. R. 2019. The histone chaperone CAF-1 cooperates with the DNA methyltransferases to maintain Cd4 silencing in cytotoxic T cells. Genes Dev, 33, 669–683.

Paul, F., Arkin, Y., Giladi, A., Jaitin, D. A., Kenigsberg, E., Keren-Shaul, H., Winter, D., Lara-Astiaso, D., Gury, M., Weiner, A., David, E., Cohen, N., Lauridsen, F. K. B., Haas, S., Schlitzer, A., Mildner, A., Ginhoux, F., Jung, S., Trumpp, A., Porse, B. T., Tanay, A. & Amit, I. 2016. Transcriptional Heterogeneity and Lineage Commitment in Myeloid Progenitors. Cell, 164, 325.

Pundhir, S., Bratt Lauridsen, F. K., Schuster, M. B., Jakobsen, J. S., Ge, Y., Schoof, E. M., Rapin, N., Waage, J., Hasemann, M. S. & Porse, B. T. 2018. Enhancer and Transcription Factor Dynamics during Myeloid Differentiation Reveal an Early Differentiation Block in Cebpa null Progenitors. Cell Rep, 23, 2744–2757.

Robinson, M. D., Mccarthy, D. J. & Smyth, G. K. 2010. edgeR: a Bioconductor package for differential expression analysis of digital gene expression data. Bioinformatics, 26, 139–40.

Rosenbauer, F. & Tenen, D. G. 2007. Transcription factors in myeloid development: balancing differentiation with transformation. Nat Rev Immunol, 7, 105–17.

Smith, S. & Stillman, B. 1989. Purification and characterization of CAF-I, a human cell factor required for chromatin assembly during DNA replication in vitro. Cell, 58, 15–25.

Subramanian, A., Tamayo, P., Mootha, V. K., Mukherjee, S., Ebert, B. L., Gillette, M. A., Paulovich, A., Pomeroy, S. L., Golub, T. R., Lander, E. S. & Mesirov, J. P. 2005. Gene set enrichment analysis: a knowledge-based approach for interpreting genome-wide expression profiles. Proc Natl Acad Sci U S A, 102, 15545–50.

Suico, M. A., Shuto, T. & Kai, H. 2017. Roles and regulations of the ETS transcription factor ELF4/MEF. J Mol Cell Biol, 9, 168–177.

Sykes, D. B., Kfoury, Y. S., Mercier, F. E., Wawer, M. J., Law, J. M., Haynes, M. K., Lewis, T. A., Schajnovitz, A., Jain, E., Lee, D., Meyer, H., Pierce, K. A., Tolliday, N. J., Waller, A., Ferrara, S. J., Eheim, A. L., Stoeckigt, D., Maxcy, K. L., Cobert, J. M., Bachand, J., Szekely, B. A., Mukherjee, S., Sklar, L. A., Kotz, J. D., Clish, C. B., Sadreyev, R. I., Clemons, P. A., Janzer, A., Schreiber, S. L. & Scadden, D. T. 2016. Inhibition of Dihydroorotate Dehydrogenase Overcomes Differentiation Blockade in Acute Myeloid Leukemia. Cell, 167, 171–186 e15.

Volk, A., Liang, K., Suraneni, P., Li, X., Zhao, J., Bulic, M., Marshall, S., Pulakanti, K., Malinge, S., Taub, J., Ge, Y., Rao, S., Bartom, E., Shilatifard, A. & Crispino, J. D. 2018. A CHAF1B-Dependent Molecular Switch in Hematopoiesis and Leukemia Pathogenesis. Cancer Cell, 34, 707–723 e7.

Wang, G. G., Calvo, K. R., Pasillas, M. P., Sykes, D. B., Hacker, H. & Kamps, M. P. 2006. Quantitative production of macrophages or neutrophils ex vivo using conditional Hoxb8. Nat Methods, 3, 287–93.

Xie, H., Ye, M., Feng, R. & Graf, T. 2004. Stepwise reprogramming of B cells into macrophages. Cell, 117, 663–76.

Zhang, Y., Gao, S., Xia, J. & Liu, F. 2018. Hematopoietic Hierarchy - An Updated Roadmap. Trends Cell Biol, 28, 976–986.

